# Large-scale investigation of species-specific orphan genes in the human gut microbiome elucidates their evolutionary origins

**DOI:** 10.1101/2023.07.17.549306

**Authors:** Nikolaos Vakirlis, Anne Kupczok

## Abstract

Species-specific genes, also known as orphans, are ubiquitous across life’s domains. In prokaryotes, species-specific orphan genes (SSOGs) are mostly thought to originate in external elements such as viruses, followed by horizontal gene transfer, while the scenario of native origination, through rapid divergence or de novo, is mostly dismissed. However, quantitative evidence supporting either scenario is lacking. Here, we systematically analyzed genomes from 4,644 human gut microbiome species and identified over 600,000 unique SSOGs, representing an average of 2.6% of a given species’ pangenome. These sequences are mostly rare within each species yet show signs of purifying selection. Overall, SSOGs use optimal codons less frequently, and their proteins are more disordered than those of conserved genes (i.e., non-SSOGs). Importantly, across species, the GC content of SSOGs closely matches that of conserved ones. In contrast, the ∼5% of SSOGs that share similarity to known viral sequences have distinct characteristics, including lower GC content. Thus, SSOGs with similarity to viruses differ from the remaining SSOGs, contrasting an external origination scenario for most SSOGs. By examining the orthologous genomic region in closely related species, we show that a small subset of SSOGs likely evolved natively de novo and find that these genes also differ in their properties from the remaining SSOGs. Our results challenge the notion that external elements are the dominant source of prokaryotic genetic novelty and will enable future studies into the biological role and relevance of species-specific genes in the human gut.

## Introduction

Genomes of the same prokaryotic species can vary substantially in gene content, giving rise to huge pangenomes, i.e., all genes present in a species(Brockhurst et al. 2019). Pangenomes generally consist of three types of genes: a small set of “core” or nearly universal genes, a larger set of “shell” or moderately conserved genes, and a huge “cloud” of rare genes(Koonin & Wolf 2008; Koonin et al. 2021). Cloud genes are particularly intriguing, as each newly sequenced genome is adding novel genes to the pangenome. This suggests that there is an ongoing appearance of genes in pangenomes, which has been mainly attributed to horizontal gene transfer (HGT). Bacteria are constantly under selection pressure to adapt to changing conditions or to colonize new niches and cloud genes might continuously provide novel genetic material on which selection can act. Thus, although most cloud genes are transient, some might prove to be adaptive and persist in the population(Conrad et al. 2021).

Pangenomes even contain genes that have no sequence similarity outside the species. Genes without detectable similarity to genes outside of a particular group have been termed lineage-specific genes, taxonomically restricted genes, or orphan genes (short: orphans)(Dujon 1996; Tautz & Domazet-Lošo 2011; Karlowski et al. 2023), and here we refer to them as species-specific orphan genes (SSOGs). SSOGs cannot be explained by the insufficient sequencing of homologous sequences, since each genome typically contains additional genes that cannot be found in other genomes, as for example observed in *Escherichia coli*(Yu & Stoltzfus 2012). Also genes specific to *E. coli*, have been found to be narrowly distributed within the species(Yu & Stoltzfus 2012); thus, most SSOGs are cloud genes and might be important contributors to the appearance of novel genetic material in pangenomes. An understanding of SSOGs could thus provide insights into cloud genes and generally into the evolutionary dynamics that shape pangenomes, contributing to an ongoing debate(Baumdicker & Kupczok 2023).

Species-specific genes have mostly been studied in eukaryotes, where they can be associated with organismal novelties and species-specific traits(Light et al. 2014), and can be important for adaptation(Khalturin et al. 2009; Santos et al. 2017). Research in eukaryotes is starting to paint a coherent picture of the evolutionary origins of SSOGs(Tautz & Domazet-Lošo 2011; Andersson et al. 2015; Prabh & Rödelsperger 2019). We now know that they can arise entirely “de novo”, from genomic sequences that are non-coding/non-genic(McLysaght & Guerzoni 2015; Oss & Carvunis 2019; Bornberg-Bauer et al. 2021) or from extensive sequence divergence beyond recognition(Vakirlis, Carvunis, et al. 2020; Weisman et al. 2020). SSOGs can also result from a combination of divergence, de novo emergence, and reuse of parts of existing protein-coding genes in alternative reading frames(Prabh & Rödelsperger 2019; McLysaght & Hurst 2016).

While most of our understanding about SSOGs and the mechanisms that give rise to them comes from eukaryotes, much less is known about them in prokaryotes. Frequently termed “ORFans”, prokaryotic SSOGs share some of the same properties as those in eukaryotes, such as shorter length and higher AT content(Daubin & Ochman 2004b). The dynamic nature of prokaryotic genomes and the known pervasiveness of HGT create additional opportunities for SSOG evolution. One hypothesis is that SSOGs initially evolve in phages or other “selfish” genetic elements and are then transferred to bacteria(Daubin & Ochman 2004b) (Figure 1). While it has been 20 years since this hypothesis was proposed, evidence to support it is limited(Daubin & Ochman 2004a; Cortez et al. 2009) and other studies failed to find equally convincing traces of the viral origin of SSOGs(Yin & Fischer 2006; Yomtovian et al. 2010).

**Figure 1:**
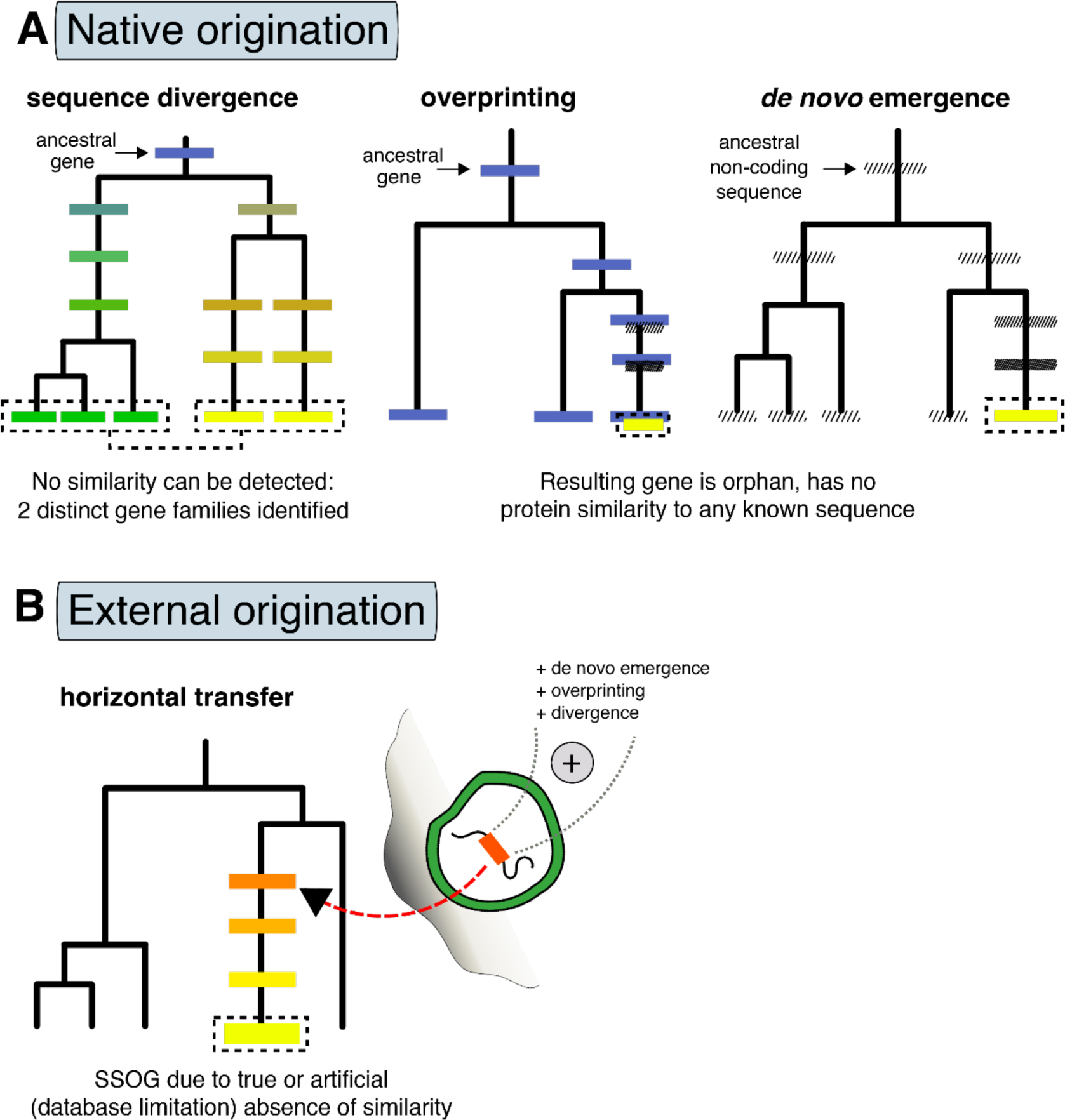
Evolutionary routes to SSOGs in prokaryotes. **A**: Native origination through sequence divergence, as well as de novo emergence from entirely non-coding regions (known as de novo emergence) or from coding regions in alternative reading-frames (known as overprinting). Overprinting can also be succeeded by gene duplication, such that both genes are then encoded in different regions. Darker shading represents transition from non-coding state to a coding one. Different colors represent differences in sequence similarity between homologous genes or in the same gene over time. **B:** Origination in phages, plasmids or integrative elements by various mechanisms such as de novo birth, overprinting, or divergence, and subsequent transfer to prokaryotic genomes. Either due to technical limitations or rapid divergence, similarity to the source of such externally originated genes might be lost, which would result in them being perceived as species-specific. Additionally, remodeling of existing genes, often in combination with overprinting and sequence divergence, can lead to SSOGs (not shown here).

An alternative explanation to SSOGs being of foreign origin is that they are of native origin(Daubin & Ochman 2004b). As for eukaryotic SSOGs, rapid coding sequence divergence, frameshifting mutations, rearrangements or a combination of the above could explain some of the genes lacking similarity, including entirely species-specific ones(Yu & Stoltzfus 2012; Lobb et al. 2015). Additionally, SSOGs could emerge de novo either from non-genic regions or from already coding ones but on an alternative reading frame, the latter known as overprinting(Delaye et al. 2008; Ardern 2023). As in eukaryotes, pervasive transcription and translation occurs in prokaryotes and may generate raw material that natural selection can shape into a functional protein(Smith et al. 2022; Wade & Grainger 2014; Wacholder et al. 2023). Artificial random peptides have been shown to readily evolve rudimentary functionality after a few rounds of selection(Knopp et al. 2019, 2021; Tenson et al. 1997), highlighting the potential of functional de novo genes to evolve in natural prokaryotic populations as well. Compared to eukaryotes, the constrained genome size, and the limited non-genic space of prokaryotes impose different constraints(Kirchberger et al. 2020). Nevertheless, there is recent support for de novo gene origination in prokaryotes: An analysis of taxonomically restricted genes from the genus *Bacillus* could identify homologous, noncoding regions in a genome of another genus for almost one third of them, supporting abundant de novo origination(Karlowski et al. 2023). Furthermore, a recent study of *Escherichia coli* showed that “remodeling events” – i.e., fusions of alternative reading-frames of segments of existing genes – are a potent source of entirely new protein sequences(Watson et al. 2022).

Finally, SSOGs can also result from annotation errors, since they are typically detected by comparative genomics approaches. These in turn rely on the automatic prediction of protein-coding open reading frames (ORFs) and sequence similarity searches between species; both potential sources of artefacts in the form of spurious ORFs and missed similarities.

To confidently assign a gene as species-specific, genome data on related species must be abundant. Recent sequencing efforts, in particular using metagenomics, where the whole community of a sample is sequenced, now provide a dense coverage of the sequence space(Chibani et al. 2022; Nayfach, Roux, et al. 2021; Coelho et al. 2022; Pavlopoulos et al. 2023). These advancements also highlighted that many genes fall into few large protein families and that there is a long tail of singleton families that have low diversity(Coelho et al. 2022). Furthermore, metagenomic data has been used to detect novel protein families with yet unknown functions that occur in multiple species(Rodríguez del Río et al. 2024). The function of a large fraction of the microbial protein universe remains unknown, where unknown sequences are often species-specific(Vanni et al. 2022), thus abundant novelty can be discovered among them.

One particularly well sampled environment is the human gut, where pangenomes of the residing species have already been constructed(Almeida et al. 2021). Those pangenomes contain the information whether a gene family is found in one or multiple species, allowing direct SSOG prediction. We build on this data – implementing further filtering steps to remove families with homology to other species – to identify putative SSOGs in the gut microbiome and to study them systematically. By looking for patterns in large-scale comparisons, we attempt to disentangle the evolutionary origin of SSOGs (Figure 1) to understand whether they originate mostly externally as suggested before, and to search for evidence of de novo evolution.

## Results

### Species-specific orphan genes are widespread in pangenomes of human gut prokaryotes

Our first goal was to establish a conservatively defined catalogue of prokaryotic species-specific orphan genes (SSOGs). We chose the human gut environment, an extensively studied niche harboring thousands of known prokaryotic species. Recently, a comprehensive catalogue of genomes from prokaryotes of the human gut was released, where the genomes were annotated and analyzed with a common pipeline(Almeida et al. 2021). In that study, 286,997 genomes were clustered into 4,644 species based on 95% identity over at least 30% of their length. Additionally, the Unified Human Gastrointestinal Protein (UHGP) catalog was clustered into homologous families at different sequence similarity cut-offs. We used the most loosely defined protein families as the starting point of our analysis (version 1.0 UHGP-50, built using a 50% identity cut-off) to identify homology with a high sensitivity. By applying a stringent pipeline of sequence-similarity searches, we filtered the initial dataset consisting of more than 600 million proteins grouped in 4.7 million families. We predicted 3.3 million species-specific proteins, grouped into 630,000 families (see Figure 2A and Methods), hereafter referred to as SSOGs (species-specific orphan genes). Unless stated otherwise, our analyses use one representative per cluster, as defined in the UHGP-50 catalog.

**Figure 2:**
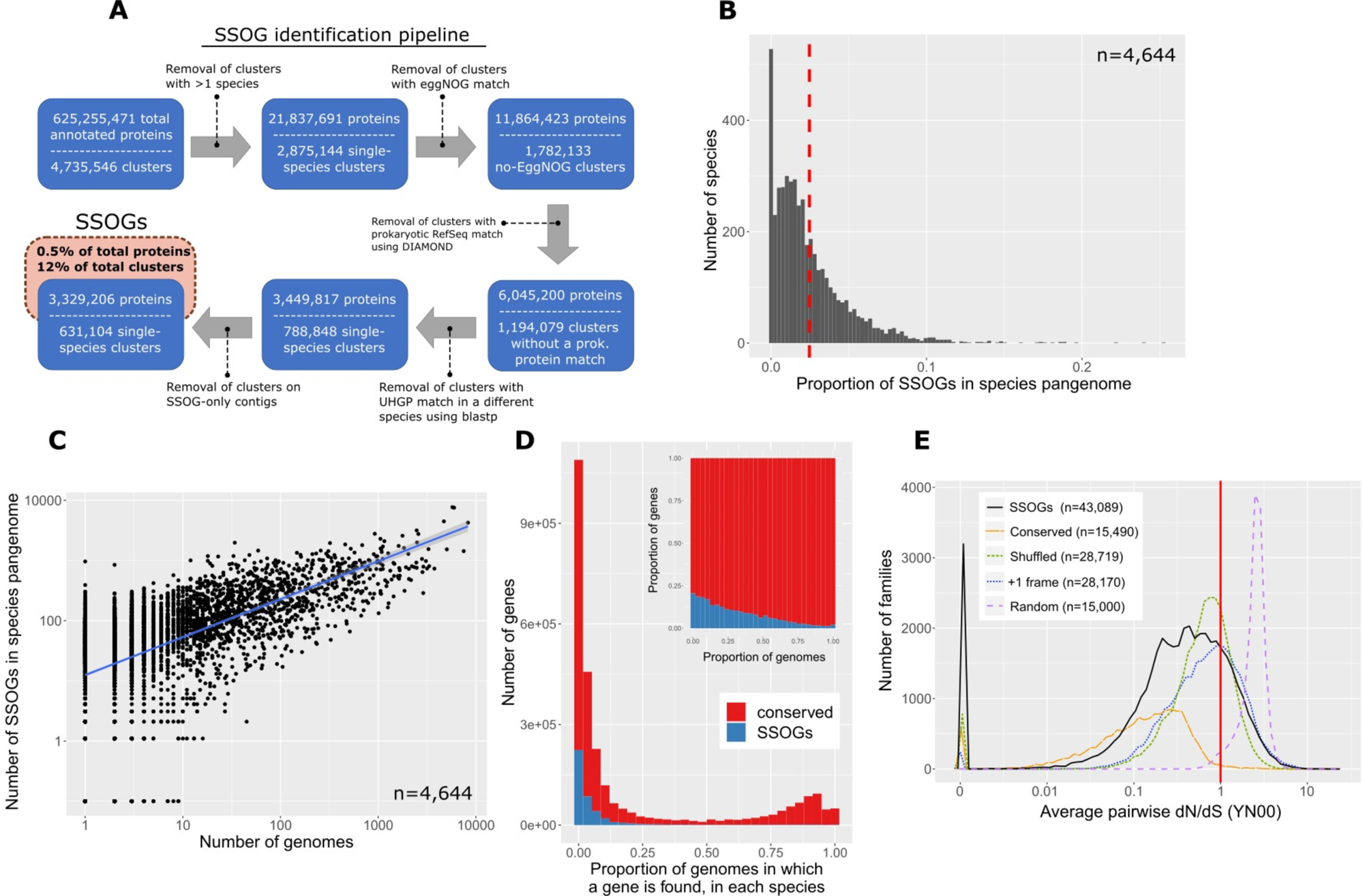
Large scale-identification of species-specific orphan genes (SSOGs) in the human gut microbiome. **A:** The main steps of the computational workflow of similarity searches. **B**: Distribution of the SSOG proportion in a given species’ pangenome. Red dashed line denotes distribution mean. **C**: SSOG number strongly correlates with the number of genomes available for a given species. **D**: Distribution of percentage of genomes where a gene is present, for SSOGs and conserved genes. Only species with at least 10 genomes are included (n=1,369). The inset plot shows the same data but as relative proportion in each bin. **E:** Distribution (60 bins) of average pairwise omega (dN/dS) per family for SSOGs (solid black line), intraspecific alignments of conserved genes (orange) and three negative controls (see Methods for details). Red line marks dN/dS=1 which corresponds to neutral evolution.

We find an average of 4.7 SSOGs per genome, and an average of 135 SSOGs per species. The latter represents an average of 2.6% of a given species’ pangenome, that is, the entire gene repertoire of the species (Figure 2B). Interestingly, the data set contains only 28 Archaea species and they harbor almost twice as many SSOGs as bacterial ones: on average 224 SSOGs per species (6.5%) and 10.9 SSOGs per genome. Notably, the underrepresentation of Archaea in sequence databases could lead to the artefactual identification of species-specific genes, since closely related species have not been sampled yet.

For all species, the number of SSOGs in a given pangenome correlates strongly with the number of genomes available for that species within the entire genome dataset provided by Almeida *et al*. (Spearman’s Rho=0.61, P<2.2*10^−16^; Figure 2C), as well as with pangenome size (Rho=0.66, P<2.2*10^−16^, partial correlation controlling for number of genomes Rho=0.39, P<2.2*10^−16^ Supp. Figure 1A), suggesting that each genome introduces novel SSOGs into the pangenome. Indeed, we find that SSOGs are rare within their respective species. Most of them are shared by at most 10% of genomes, i.e., belonging to the cloud fraction of prokaryotic pangenomes (Figure 2D). When compared to conserved genes, i.e., all non-SSOGs, SSOGs are strongly overrepresented among accessory genes (Supp. Figure 2). While this has been reported before in *E. coli*(Yu & Stoltzfus 2012), our analysis demonstrates that this is a general feature of the human gut microbiome and potentially of prokaryotic SSOGs in general. Thus, SSOGs are indeed widespread and we find them in in 94% of all gut species and in all species with at least 10 available genomes. Nevertheless, they represent only a small fraction of the gene repertoire of a particular species and they tend to be rare within pangenomes.

We observe variability across taxonomic groups: some bacterial taxonomic classes have markedly higher SSOG percentages than others, with Coriobacteriia representing the lower (avg. 0.7%) and Alphaproteobacteria the higher (avg. 5.4%) extremes (see also Table 1 and Supp. Figure 3A). Next, we investigated the impact of metagenome assembled genomes (MAGs) on SSOG numbers. Species with a high proportion of MAGs have comparable SSOG percentages as species with many isolate genomes (Supp. Figure 1B), and species with lower quality assemblies show no trend towards more SSOGs (Rho=-0.04, P=0.0024; Supp. Figure 1C). This suggests that the estimated SSOG numbers are not driven by assembly artefacts.

**Table 1:** Basic statistics of SSOGs and their various subsets for the nine most represented taxonomic classes.

| Taxonomic class | No. of species | No. of SSOs | Pct. of SSOs out of species pangenome (average) | No. of SSOs dN/dS < 0.5 | No. of SSOs >800nt | Pct. of SSOs that are cloud | No. of SSOs with viral hits | No. of SSOs matching alt. frames of conserved genes | No. de novo candidates |
| --- | --- | --- | --- | --- | --- | --- | --- | --- | --- |
| Actinobacteria | 124 | 18932 | 4.06 | 535 | 472 | 85 | 268 | 261 | 40 |
| Alphaproteobacteria | 61 | 13398 | 5.38 | 569 | 721 | 79.6 | 174 | 252 | 15 |
| Bacilli | 488 | 40630 | 2.20 | 2311 | 1295 | 81.3 | 3598 | 400 | 56 |
| Bacteroidia | 603 | 189793 | 4.41 | 5576 | 2954 | 87.8 | 4074 | 3068 | 389 |
| Clostridia | 1868 | 218177 | 2.38 | 9760 | 10098 | 86 | 9667 | 2390 | 317 |
| Coriobacteriia | 704 | 21808 | 0.72 | 976 | 673 | 91.4 | 517 | 204 | 41 |
| Gammaproteobacteria | 288 | 45025 | 2.51 | 1497 | 1340 | 89.7 | 1151 | 743 | 127 |
| Negativicutes | 148 | 18916 | 2.12 | 922 | 437 | 90.6 | 1716 | 317 | 11 |
| Vampirovibrionia | 63 | 5794 | 2.34 | 363 | 178 | 77.2 | 542 | 66 | 11 |
| all | 4644 | 631104 | 2.6 | 25355 | 21493 | 86.8 | 23147 | 8477 | 1075 |

Genes that show protein-coding conservation typically exhibit a higher rate of synonymous polymorphisms (dS) compared to nonsynonymous polymorphisms (dN), i.e., dN/dS<1. In contrast, this pattern is not expected for spurious open reading frames (ORFs), i.e., ORFs incorrectly annotated as protein-coding. The SSOGs predicted here lack homologues in other species (by definition) and tend to be rare within their respective species, which could be an indication that they are spurious. To test the impact of spurious ORF prediction on our SSOG dataset, we estimated dN/dS from intraspecific multiple sequence alignments for each of the 154,650 SSOG families with at least two non-identical sequences. Due to the high degree of genetic similarity, a dN/dS ratio could be obtained for only 43,089 of them (see Methods). The distribution of values is clearly shifted to the left of 1 with a median of 0.38 and a mean of

0.63 (Fig. 2E) suggesting that a considerable proportion of these genes are under selection. To provide further evidence, we calculated dN/dS in the same manner for three different negative noncoding controls (see Methods). Crucially, the dN/dS distribution of SSOGs is distinct from all three controls (all Wilcoxon test P-values < 2.2*10^−16^), has a second peak of values below 1 that is missing from controls and has significantly more genes without any nonsynonymous mutations (that is with dN/dS = 0; 7.4% vs. 2.7%, 0.9% and 0%). Importantly, our findings are robust to the choice of methodology, as we obtained similar results when estimating selection using a different approach and tool (see Methods and Supp. Figure 3B) as well as when only including genes over 300nt in the analysis (n=21,203 leaving out half of our dataset; Supp. Figure 3C). As a positive control, we also calculated dN/dS on intraspecific alignments of conserved genes in 9 randomly selected species (see Methods; Fig 2E). This distribution is closer to 0 (median 0.14) but extensively overlaps that of SSOGs, and to a lesser degree that of noncoding controls. In other words, SSOGs lie somewhere between conserved and noncoding controls, but with large overlaps.

The landscape of these dN/dS values should be interpreted with two important points in mind. First, even when selection is present, the power to detect it is limited within the same species(Kryazhimskiy & Plotkin 2008). Second, young, newly evolved genes can be functional even when they show no evidence of selection(Wacholder et al. 2023; Vakirlis et al. 2022). Thus while we cannot conclude that all SSOGs with a dN/dS close to 1 are not functional genes, this analysis strongly supports that, at the very least, a considerable SSOG proportion is under selection, and thus functional. Based on the above analysis, we define a high-confidence set of SSOGs with a dN/dS < 0.5 (n=25,355).

### SSOGs and their proteins have some distinct and some common properties compared to conserved genes

In both prokaryotes and eukaryotes, species-specific genes are known to exhibit certain characteristics, such as shorter length, which set them apart from conserved genes(Tautz & Domazet-Lošo 2011; Yomtovian et al. 2010; Vakirlis et al. 2018). These gene and protein properties might provide clues about the evolutionary origins of SSOGs. To compare SSOGs to conserved genes, we next turned to properties of genes and proteins across species.

Across every taxonomic class, SSOGs are significantly shorter than conserved genes, at about a third of their average length (Figure 3A), and this is true also for the high-confidence set (Supp. Figure 4A) There is only a weak correlation between the average length of SSOGs and conserved genes per species (Rho=0.14, P<2.2*10^−16^). Average GC content of SSOGs in a given species is strikingly similar (avg. 0.45 vs. 0.46) to that of conserved genes (Figure 3B), exhibiting an almost one-to-one correlation (Rho=0.96, P<2.2*10^−16^). This also holds for the high-confidence set (same averages; Rho=0.92, P<2.2*10^−16^; Supp. Figure 4B) and when only considering SSOGs longer than 800nt (Supp. Figure 4C) demonstrating that this similarity is not explained by the existence of spurious ORFs. We see the same trend when restricting the analysis to Archaea (Supp. Figure 4D), as well as when comparing SSOGs and conserved genes within individual species (Supp. Figure 4E). The trend also applies to GC content in the 3^rd^ synonymous position of codons (GC3s; see Supp. Figure 4F) and for CpG content (Supp. Figure 5). This observation fits the native origination scenario, in which SSOG GC content simply reflects the genome’s GC content, but, in theory at least, it would also be consistent with external transfer followed by rapid adaptation of the incoming gene’s GC content to that of the host genome(Lawrence & Ochman 1997).

**Figure 3:**
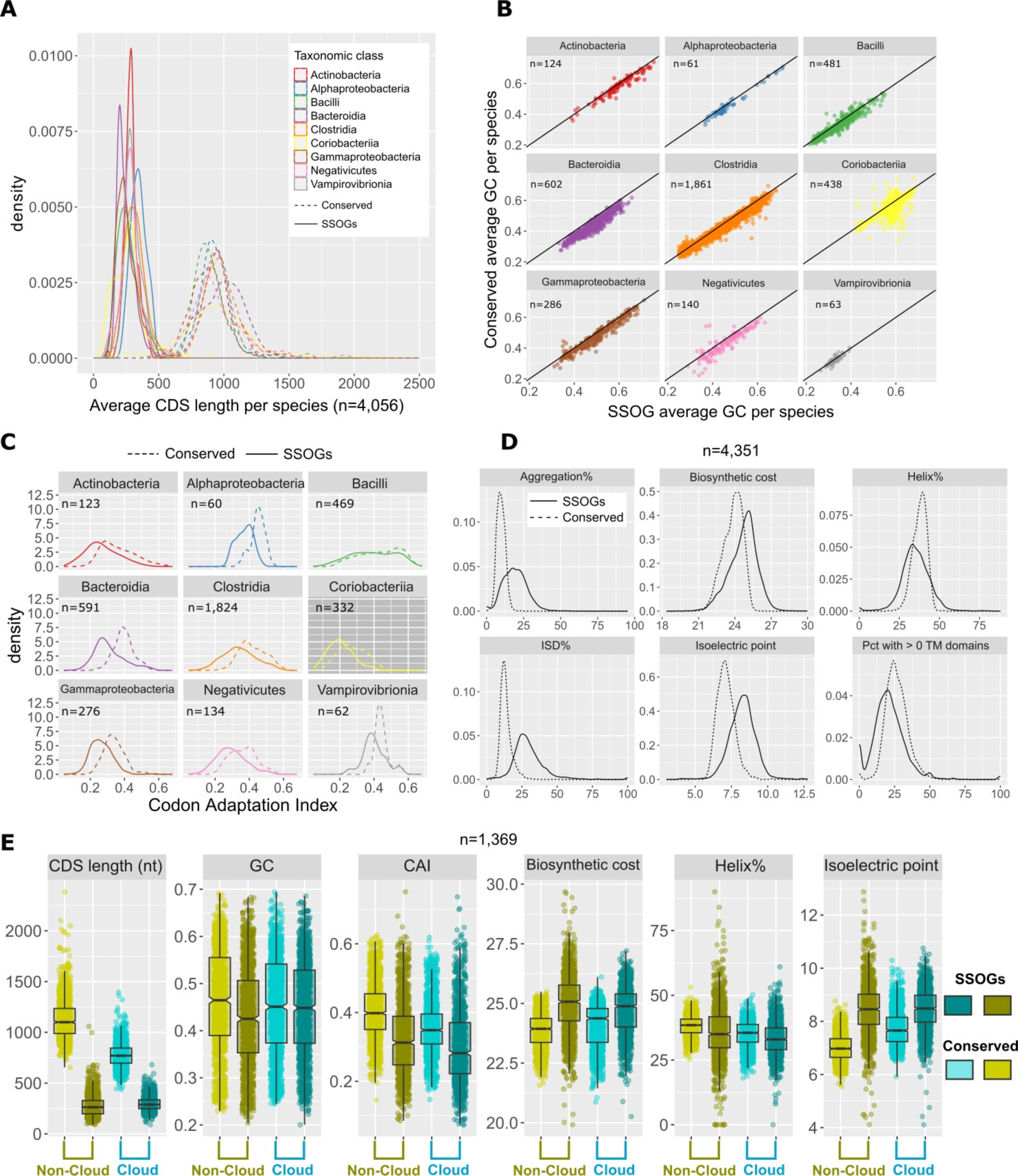
Properties of SSOGs compared to conserved genes. **A:** Density plots of the average length of SSOGs and conserved CDSs in each species, grouped by each of the 9 best represented taxonomic classes of our dataset. Only classes with at least 50 species are included. **B**: Average GC content of SSOGs and conserved genes in each species, grouped by taxonomic class (same classes as in A). **C**: Same comparison as in A (average per species), but with Codon Adaptation Index (CAI) values as predicted by the *CAIJava* tool. All Wilcoxon test P-values < 10^−4^. **D**: Comparison of proteins encoded by conserved genes and SSOGs, in terms of average percentage of protein predicted to self-aggregate, biosynthetic cost, percentage of protein predicted to be helical, percentage of protein predicted to be disordered (ISD), isoelectric point of a protein, and percentage of proteins with at least 1 transmembrane domain. All Wilcoxon test P-values < 2.2*10^−16^. **E**: Comparison of some of the properties found in panels A-D in 1,369 species with at least 10 genomes where SSOG and conserved gene categories are split in cloud (present in 10% of genomes or less) and non-cloud. All plots show points or distributions of average values per species.

In certain origination scenarios, including de novo evolution and recent transfer from external origin, SSOGs are expected to show suboptimal codon usage relative to the rest of the genome. We thus calculated the Codon Adaptation Index (CAI), a measure of how frequently optimal codons (preferred at the level of an entire genome) are used by each gene, for all genes of the representative genome of each species using the *CAIJava* tool, (which does not require a reference gene set) and compared CAI of SSOGs and conserved genes (see Methods). Across taxonomic classes, SSOGs have consistently lower CAI values than conserved genes (Figure 3C, all Wilcoxon test P-value < 10^−4^). This is also true when controlling for the length differential between SSOGs and conserved genes by subsampling conserved genes to obtain a set of comparable mean length to that of SSOGs (Supp. Figure 6A; Methods). It is also true for both the high-confidence set and for the set of SSOGs longer than 800nt (Supp. Figure 6B and 6C). For *E. coli*, we also calculated CAI based on codons used in reference sets of highly expressed genes and found similar results (Supp. Figure 7; Methods).

At the protein level too, SSOGs exhibit distinguishing characteristics (Figure 3D). Compared to conserved genes, they are, on average, richer in intrinsically disordered regions, a smaller percentage of their sequence is found in helical regions, a smaller percentage of them contains transmembrane (TM) domains, they are more aggregation prone, have higher biosynthetic cost, and a higher isoelectric point (the latter was also observed for de novo genes in yeast and fly(Montañés et al. 2023; Blevins et al. 2021)). All these differences hold when considering each taxonomic class individually (Supp. Figure 8). However, when controlling for the length differential between SSOGs and conserved genes, intrinsic disorder and TM domains are no longer significantly different (Supp. Figure 9). Note that novel genes in membrane-related roles have previously been suggested for eukaryotes such as budding yeast(Vakirlis, Acar, et al. 2020; Tassios et al. 2023)and for prokaryotes(Sberro et al. 2019). Here we find that, even though SSOGs have similar nucleotide composition to conserved genes on average (and even more so when controlling for length differences) they differ in key properties, such as their length, codon usage, and some protein structural features.

Since we observed before that most SSOGs are cloud genes, i.e., rare in the species where they are found (Figure 2D), we next asked if cloud SSOGs differ from non-cloud SSOGs. Indeed, we find that some properties differ, e.g., cloud SSOGs have lower CAI, higher GC content, and fewer helices than remaining SSOGs (Figure 3E). Surprisingly, when comparing cloud genes to remaining genes, the observed trend for GC content differs between SSOGs and conserved genes: whereas conserved cloud genes have a lower GC content than remaining conserved genes, SSOG cloud genes have higher GC content than the remaining cloud genes. Furthermore, the GC content of SSOG cloud genes is higher than the GC content of conserved cloud genes. These trends also hold when analyzing taxonomic classes separately (Supp. Figure 10) and when zooming in on individual species (Supp. Figure 4C).

Cloud genes could be explained by a continuous inflow of genes by horizontal gene transfer, which is expected to result in conserved cloud genes. We believe that SSOG cloud genes have different evolutionary origins than conserved cloud genes due to their striking difference in the trends for GC content. Interestingly, SSOG non-cloud genes have low GC content, which can be explained by de novo origination from noncoding regions which typically have lower GC content than coding regions or by rapid divergence, which could lead to the accumulation of A and T(Hershberg & Petrov 2010). Rapid divergence and transcriptional silencing by the histone-like nucleoid structuring protein (H-NS) has been described for genes of low GC content in Salmonella(Papanikolaou et al. 2009)which might also apply to the SSOGs with low GC content.

### SSOGs of likely phage origin have distinct characteristics

Here, SSOGs are defined as species-specific genes with respect to other prokaryotes, however it is possible that they are the result of recent horizontal gene transfer from bacteriophages. To identify such transferred genes, we conducted similarity searches of the SSOGs against two recently published phage protein databases, the metagenomic gut virus catalog (MGV)(Nayfach, Páez-Espino, et al. 2021) and the gut phage database (GPD)(Camarillo-Guerrero et al. 2021), that are partially overlapping but mostly complementary in terms of protein sequence content.

We find that SSOGs are strongly depleted in similarity matches to phage proteins relative to conserved genes, ranging from 3 to 6-fold using different parameters and cut-offs (Figure 4A; Methods). At low stringency, 5.1% of SSOGs have similarity to viral sequences, compared to 63% of conserved genes. SSOGs are also depleted, when separately searching against either MGV or GPD (Supp. Figure 11A), when using the high-confidence set and when excluding short sequences (<800nt) for which similarity searches often fail (Supp. Figure 11B and 11C). Interestingly, binning conserved genes according to the number of species in which they are found (a proxy of their evolutionary age) reveals a pattern that the proportion of genes with similarity to phages increases with the number of species where the gene is found (Figure 4B), regardless of the stringency criteria applied (Supp. Figure 11D). This pattern is inconsistent with a scenario under which most SSOGs arrive through HGT from bacteriophages, which we would expect to either produce the inverse trend, if only a fraction of HGT is ultimately retained, or no trend at all in the case of a constant rate of transfer and retention.

**Figure 4:**
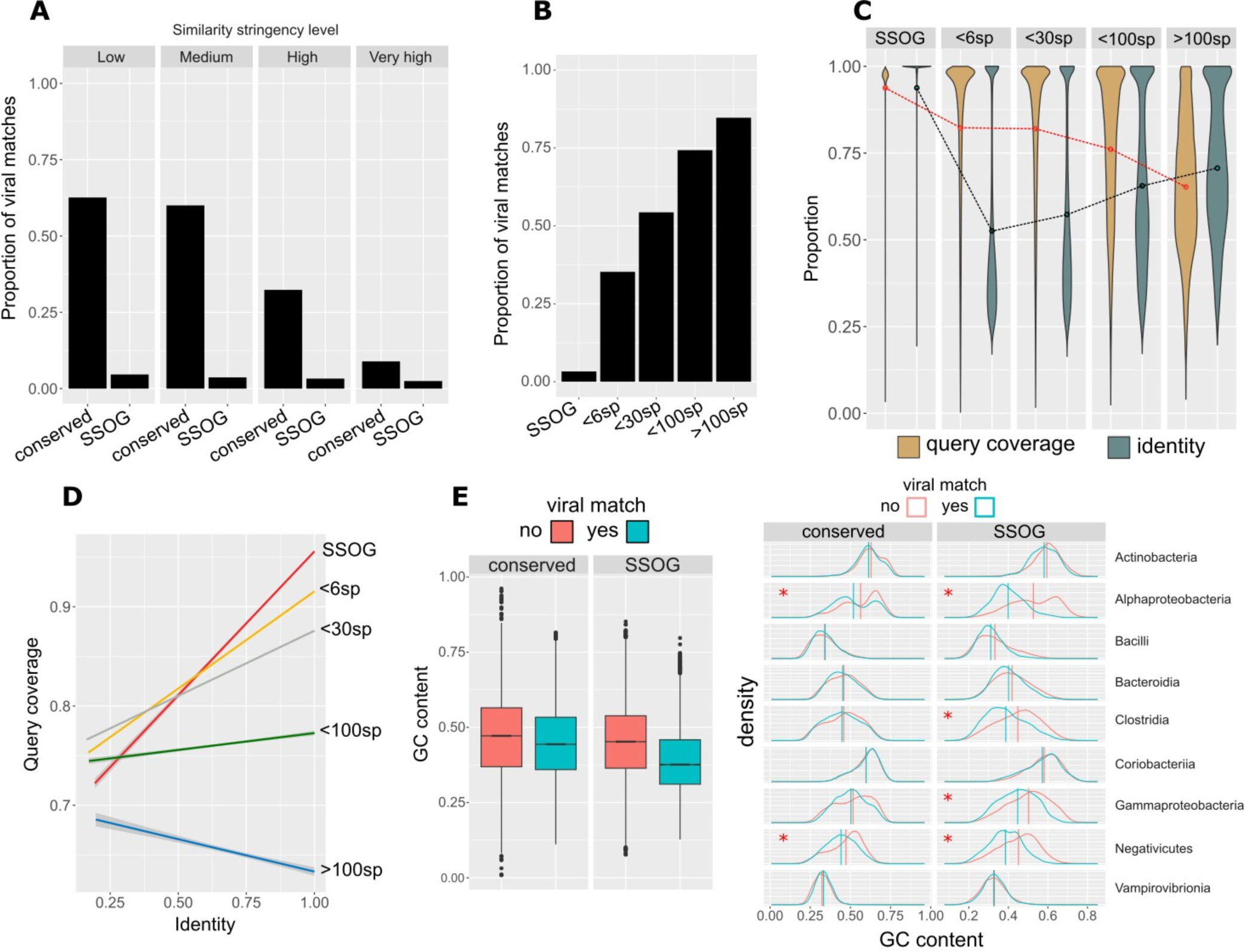
Properties of genes with viral matches. **A**: Proportions of conserved proteins and SSOGs with statistically significant similarity matches to viral proteins at four different significance thresholds. **B**: Proportions of proteins with statistically significant similarity matches to viral proteins (at high stringency level), binned according to the number of prokaryotic species (sp.) in which a homologue can been found (size of the prokaryotic protein family). **C**: Query (prokaryotic protein) coverage and sequence identity of top statistically significant matches (all hits with E-value < 10^−5^ are included), binned according to the number of species in which a homologue can be found. Lines connect distribution means. **D**: Correlation between query coverage and identity in sequence matches involving genes belonging to different sizes of protein families, same data as in C. **E**: Distributions of GC content of genes with and without a statistically significant viral match (high stringency), among all SSOGs and conserved genes (boxplot) and among the nine best represented taxonomic classes (density plots). A red asterisk denotes a non-negligible effect size in difference of means calculated by Cliff’s Delta (Delta estimate > 0.15). Note that in Alphaproteobacteria and Negativicutes the difference exists also in conserved, but the effect size there is much weaker compared to SSOGs (−0.21 vs. −0.62 and −0.2 vs. −0.42).

As both GPD and MGV databases have been assembled with the use of automated annotation pipelines, it is possible that they may contain bacterial contigs that are falsely classified as coming from phages. To control for such a bias, we confirmed our findings by searching a third phage genome database(Shah et al. 2023) which is smaller but manually curated and was built from samples taken from infant guts. We observed similar patterns as for the larger databases confirming that the proportion of SSOGs (and more generally, less conserved genes) with similarity to phage proteins is considerably lower than that of widely conserved genes (Supp. Figures 11E and 11F). The overall proportion of matches in all gene groups is lower compared to GPD and MGV, but that is expected given the difference in size (10K phage OTUs vs. 190K and 142K in MGV and GPD respectively).

The trend of increasing percentage of phage matches with increasing conservation across species prompted us to investigate further. To dissect the evolutionary dynamics, we plotted, across age categories, the query coverage and percent identity of the best phage match per protein as defined by E-value, without filtering any high-identity hits (Figure 4C). This revealed that matches to SSOGs are very close to 100% identity and coverage with both averages decreasing with age. Yet, while the coverage distribution shows a clear downward trend, the identity distribution becomes bimodal with age, with one peak staying near 100% and another one initially appearing at around 35%, then increasing. Furthermore, while identity and coverage are strongly correlated for viral hits of SSOGs, this correlation gradually decreases with age (Figure 4D). One explanation for these trends is that a subset of genes has only recently been transferred between phages and bacteria, while another subset consists of older transfers that have undergone divergence, truncation, or both.

We previously established that SSOGs mirror conserved genes’ averages in crucial aspects such as nucleotide composition. Given the evidence that a small percentage of SSOGs appears to have been recently transferred from phages, we might expect that they exhibit distinguishing signatures at the level of their nucleotide composition. Indeed, we find that overall, SSOGs with similarity to phage proteins have markedly lower GC content than those without (Figure 4E), congruent with what is known for phage GC content relative to their hosts(Almpanis et al. 2018). Most strikingly, the effect of this difference is practically negligible when looking at conserved genes, albeit staying statistically significant due to the large sample size. These results hold when only genes >800nt are considered (Supp. Figure 11G) and when using the high-confidence SSOG set (Supp. Figure 11H). We thus conclude that there is a low proportion of SSOGs with hits to phages that appear distinct from the remaining SSOGs.

Additionally, we computationally detected prophage regions in all genomes using Phigaro(Starikova et al. 2020). We find that a very low proportion of SSOGs reside within prophages (13,174/3,329,206; 0.3957%), which is slightly higher than the overall proportion of prophage genes among all genes (2,029,792/625,255,471; 0.3246%). Thus, although phage origin might play a role, this can only explain a small fraction of SSOGs.

### A subset of SSOGs shows evidence of native origination

For some SSOGs, it might be possible to trace the origin within their native genome, either to existing genes or to non-coding regions. We first searched for SSOGs that might have originated via utilization of an alternative frame following duplication or horizontal gene transfer(Tria & Martin 2021) from closely related genomes. To this end, we performed similarity searches of six-frame translations of all SSOG coding sequences against six-frame translations of all coding-sequences of their native genome. We found that only a small percentage, 1.34% of SSOGs (8,477 representative SSOGs), have at least one significant match to a conserved gene (i.e. non-SSOG) excluding gene pairs with overlap. 75% of matches are on the reverse strand of the conserved gene, suggesting a more frequent exploration of the reverse complement sequence space rather than the alternative reading frames of the same strand. Out of all SSOGs matching conserved genes, 64% (5,397) match a single conserved gene, 26% (2,238) match two conserved genes and the rest (10%) match three or more. SSOGs matching more than one conserved gene correspond to chimeras similar to what was found for *E. coli*(Watson et al. 2021).

Approximately 10% of SSOGs with hits (892) match conserved sequences on the forward strand and the same frame as the conserved gene. Thus, these should be considered as false positives, that is, non-SSOGs. Such cases can be explained by the impact of the database size on the calculation of the E-value, which is used to filter the matches. The remaining 7,585 SSOGs with matches are strong candidates for having evolved out of alternative reading frames of preexisting native proteins, although they could also be explained by annotation errors of overlapping ORFs. For SSOGs with matches on the same contig (n=1,515, 20%), the genomic distances between the SSOG and the matching gene show a unimodal distribution with a mean of 131kb (Figure 5B).

**Figure 5.**
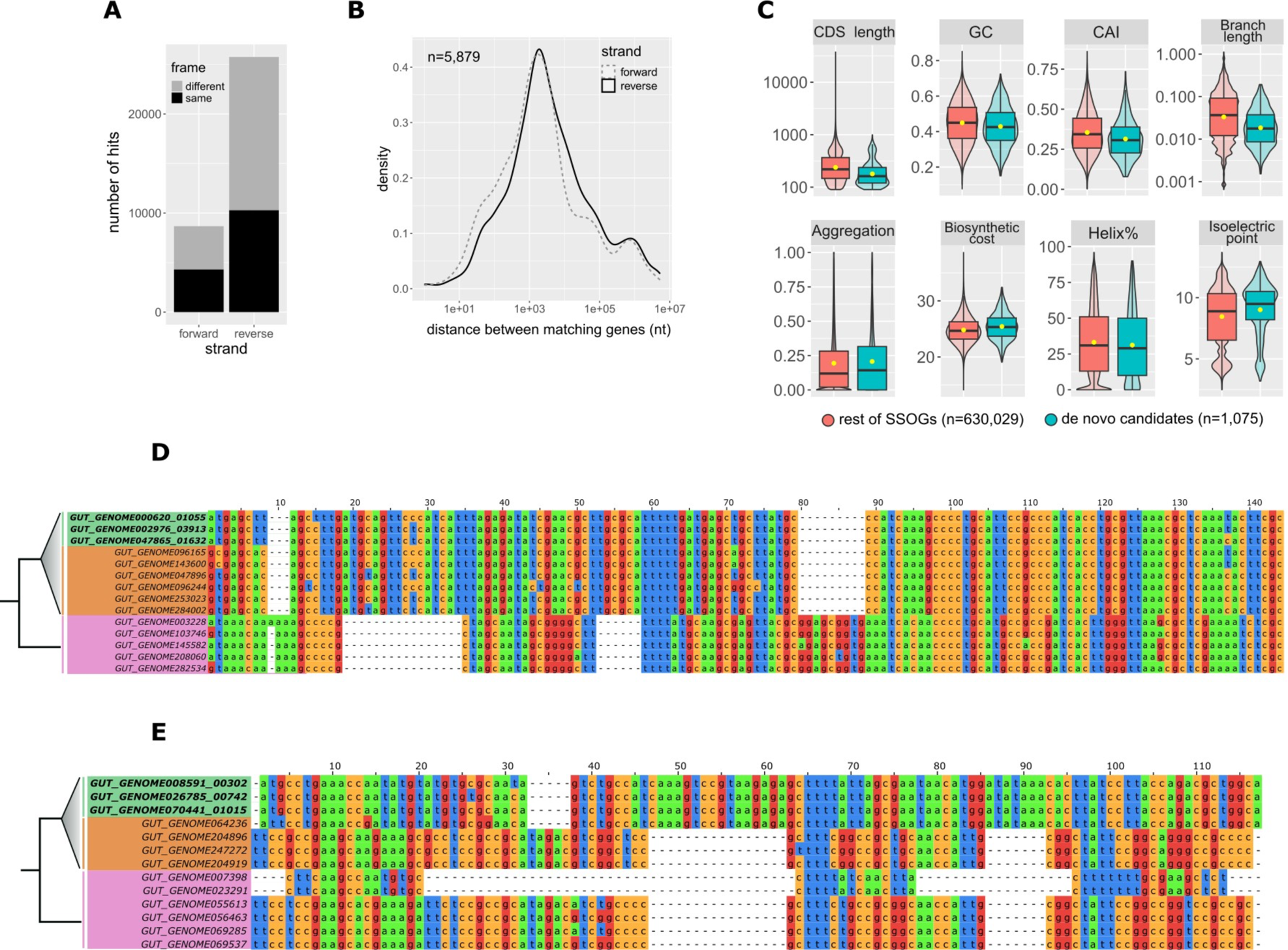
Evidence for native origination of SSOGs. **A**: distribution of frame and strand of similarity matches between SSOGs and conserved genes within each genome. **B**: Distribution of genomic distances between matching SSOG and conserved gene, when these are found on the same contig. **C**: Comparisons of de novo candidates to remaining SSOGs: Distributions of Codon Adaptation Index (CAI), terminal species-level branch length, proportion of protein sequence predicted to be aggregation-prone, biosynthetic cost, CDS length, GC content, protein percentage estimated to fold into a helix, and isoelectric point. Details of statistical comparisons can be found in the main text. Helix% comparison Wilcoxon test P-value = 0.0064. Aggregation% P-value = 0.26. Branch length and CDS length are shown on log scale for visibility. Outliers are not shown as points but are represented by the violin plot range. **D**: Alignment of a de novo candidate gene from *Hafnia paralvei* (green, three non-identical members of an eight-member family are shown) to its orthologous regions in genomes of the same species that do not have annotated homologues (orange) and genomes of its closest outgroup species (pink). Identical sequences have been removed from the alignment. The average pairwise dN/dS for this family is 2.5. **E**: Same as D, but for another candidate from an unnamed species of the order Opitutales. Three non-identical members of a family of six members are shown. Five orthologous loci (i.e. from five genomes) that contain a long insertion have been removed for visual purposes, and an additional 37 orthologous loci are not shown again for visual purposes (their sequences are almost identical to those shown in the figure). The average pairwise dN/dS for this family is 3.4. Alignments were generated with MAFFT(Katoh & Standley 2013) and visualized with JalView(Waterhouse et al. 2009).

We next turned to evidence for native origination that can be found in closely related species. A robust approach for the detection of de novo gene emergence is the identification of the orthologous genomic region in outgroup genomes. By examining such regions, it is possible to establish that a gene has evolved out of previously non-coding sequences, or out of an already coding locus, either coexisting with or replacing an ancestral gene. To detect both such cases, we identified a given SSOG’s orthologous region in outgroup genomes of the same species and those of its closest relative species using strict orthology criteria (see Methods) and discarded cases with missed orthologous open reading frames (ORFs) that might have eluded the automatic annotation pipeline. We thus identified 1,075 SSOGs that can be considered candidates for de novo emergence from non-coding regions (Supp. Table 1).

Closely related species are more likely to share genomic synteny and hence to satisfy our conservative criteria. The de novo candidates originate from 718 different species, and these have shorter terminal branches in the tree compared to the species of the remaining SSOGs (Figure 5C; means of 0.03 and 0.071 substitutions per site; Wilcoxon test P-value < 2.2*10^−16^; Cliff’s Delta 0.28). Thus, they represent the evolutionarily younger end of the spectrum of all de novo gene candidates, as expected. Comparisons of gene and protein properties support this view (Figure 5C). De novo candidates are overall shorter (Delta=0.28; P<2.2*10^−16^), have lower CAI (Delta=0.18; P=3.64*10^−11^) and GC content (Delta=0.1; P=8.5*10^−9^), have slightly higher isoelectric point (Delta=0.15; P=10^−13^) and biosynthetic cost (Delta=0.13; P<2.2*10^−16^). Since mutation is biased towards AT, most bacteria have lower GC content in noncoding regions(Hershberg & Petrov 2010), which can explain the lower GC content in de novo candidates compared to remaining SSOGs. Additionally, de novo candidates are depleted in matches to viral proteins compared to the rest of SSOGs (Fisher’s test odds-ratio: 40.9, P-value = 3.5*10^−16^). Two examples of alignments of de novo candidates to their orthologous regions in outgroup genomes can be found in Figure 5D and E. The translated sequences can be found in Supp. Figure 12, together with two additional examples and their translations (selected for visualization based on their short alignment length). In both these cases, based on the multiple stop codons present in the orthologous sequences, we can cautiously infer that the ancestral sequence did not contain an ORF and thus lacked coding potential.

### Operon-like arrangements provide functional hints for some SSOGs

Assigning a function to SSOGs is notoriously difficult. For instance, we searched a database from a previous study that established mutant phenotypes for thousands of prokaryotic genes coming from 46 strains from 36 different species(Price et al. 2018) including many from our dataset (see Methods), but found no phenotypes for any of the SSOGs.

Even if no experimentally derived functional information has been determined for a SSOG, its genomic context can still offer some clues as to what its role might be in the cell(Coelho et al. 2022; Osbourn & Field 2009). We thus examined the neighbors of all SSOGs (not only representative ones) in all genomes harboring sSOGs to identify cases of operon-like arrangements where one functional term is present in multiple neighbors (see Methods). We identified 1,905 SSOGs in operon-like arrangements, belonging to 405 distinct protein families. For these, at least 3 out of the 6 closest neighbors (i.e. 3 downstream and 3 upstream) have a common Gene Ontology (GO) term associated. We then looked for enrichment of specific GO terms within the SSOGs against the background of all genes in operon-like arrangements (n= 4,362,138). The most overrepresented specific terms by gene counts are shown in Figure 6A. The results of the full GO term enrichment analysis can be found in Supp. Table 2.

**Figure 6.**
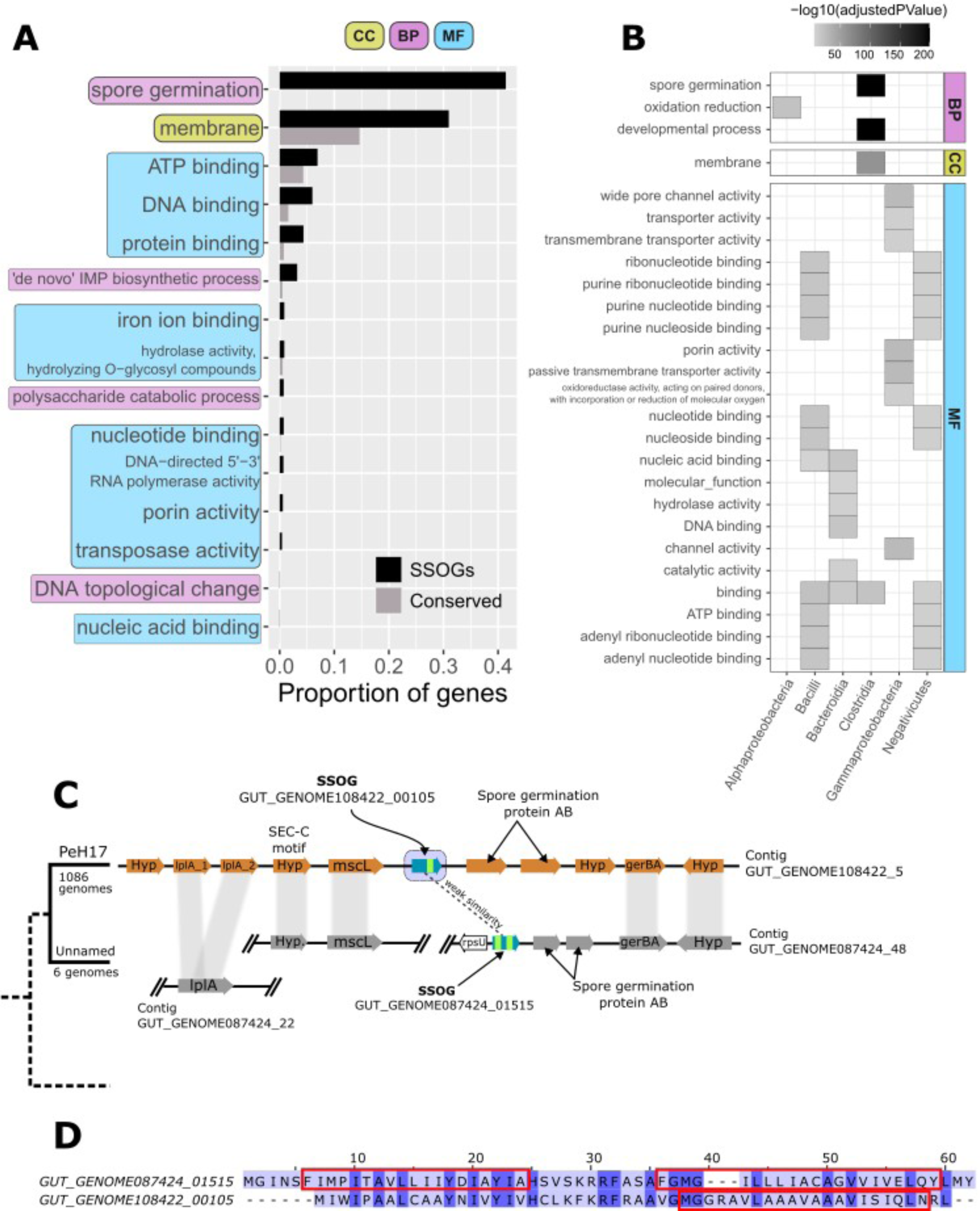
Genomic context offers functional clues of SSOGs. **A**: Proportion of SSOGs and conserved genes annotated with a given GO term based on genomic context. These GO terms are those with the highest SSOG proportion and are all significantly overrepresented with an FDR adjusted P-value < 10^−5^. The complete results of the GO enrichment analysis can be found in Supp. Table 2. CC: Cellular Component. BP: Biological Process. MF: Molecular Function. **B**: Significantly overrepresented GO terms among SSOGs, when analyzing separately genomes of species belonging to each of the nine best represented taxonomic classes. For visualization purposes, only GO terms significantly overrepresented with an FDR adjusted P-value < 10^−5^ and found in at least 10% of SSOGs in a class are shown. The complete lists, including for two of the nine classes (Actinobacteria, Vampirovibrionia) for which no enrichment was strong enough to be included in this figure, can be found in Supp. Table 3. (no enrichment was found for Coriobacteriia). **C**: Genomic region around Clostridia species PeH17 SSOGGUT_GENOME108422_00105 and syntenic conservation in its sister species (Unnamed, representative genome GENOME056772). Gene names are included in each gene shape. Vertical grey bands connect genes found within the same homologous family. Blue arrows denote SSOGs, and green bands within them represent TM domain encoding segments. lplA: lipoate-protein ligase A. mscL: Large Conductance Mechanosensitive Ion Channel protein. gerBA: spore germination protein B1. Hyp: hypothetical protein. No annotation was present in the GFF files for the two proteins to the right of the SSOGs, but all four had the same match in InterPro (Spore germination protein AB; IPR004761). **D:** Pairwise alignment generated by MAFFT, of the protein sequences of the two SSOGs highlighted in C. Rectangles mark the positions of predicted TM domains in the two protein sequences.

The term “membrane” was the only cellular component that was found in at least 3 SSOG neighbors and it was one of the top enriched terms overall (GO:0016020; Figure 6A, Supp. Table 2). This is intriguing given previous findings linking TM domains to novel genes(Vakirlis, Acar, et al. 2020; Sberro et al. 2019). A closer look showed that the membrane enrichment was due almost exclusively to SSOGs from the class Clostridia. Thus, to account for taxonomic specificities, we repeated the analysis separately for each of the 9 best represented taxonomic classes and found limited overlap between the GO terms enriched within each class (Figure 6B; Supp. Table 3).

At the level of biological processes, one of the strongest enrichments was “spore germination” and this enrichment too is specific to Clostridia (Figure 6B). Of the seven Clostridia SSOGs related to this term, four had at least one TM domain (57%) including three that are also annotated as “membrane”, which is expected given that the cell membrane is central to the process of germination. We examined in detail one large family (n=137) with “spore germination” annotation and strong selection evidence (dN/dS=0.25; see Methods). A closer look at the wider genomic region of this family showed clear evidence of syntenic and functional conservation with its closest neighboring species (unnamed species, see Figure 6C). Furthermore, another SSOG, also encoding TM domains, is found at the predicted orthologous locus in the closest species. Alignment of these two SSOG protein sequences of similar size reveals traces of similarity and an almost complete overlap of the TM domains in each sequence (Figure 6D). This intriguing finding suggests that these two genes are homologous and likely functionally similar but have diverged beyond detectable similarity, as observed before(Vakirlis, Carvunis, et al. 2020; Weisman et al. 2020). When their common ancestor might have first evolved is unclear. Overall, this analysis suggests that SSOGs might evolve different functions in different lineages and point towards spore germination as a particularly relevant process in Clostridia.

## Discussion

Species-specific genes are ubiquitous. Across the tree of life, they may be seen as one end of a taxonomic gradient, with universally conserved genes at the other end and genes restricted to smaller taxonomic groups as intermediates(Tautz & Domazet-Lošo 2011). Some recent large-scale studies have focused on novel prokaryotic gene families present in more than a single species and have firmly demonstrated their functional relevance and hinted towards crucial habitat-specific roles(Coelho et al. 2022; Sberro et al. 2019; Río et al. 2022; Pavlopoulos et al. 2023). The question of the evolutionary origin of novel prokaryotic families however was mostly left unexplored. This question is best approached by focusing on orphan genes that are species-specific and thus represent the most-recently evolved form of genetic novelty.

Here we predict species-specific orphan genes (SSOGs) in the human gut microbiome and show that these SSOGs evolved due to multiple routes. We find that a low proportion of them has matches in viral proteins. Thanks to the extensive, high-quality sampling of both the prokaryotic and the viral diversity of the human gut, we can be relatively confident that our findings are not significantly biased by limitations of the dataset. Additionally, SSOGs with viral matches have a distinct composition compared to the remaining SSOGs (Figure 4E), which is inconsistent with the external origination scenario for most SSOGs. Conversely, there is a low proportion of genes, where we can recover evidence for de novo origination from non-coding sequences. These SSOGs also differ from the remaining SSOGs by being shorter, having lower GC%, and lower CAI (Figure 5C), although these differences likely also reflect the fact that we can only detect the very youngest of the de novo originated orphans. Nonetheless, these results are also inconsistent with de novo origination from non-coding sequences as a dominant source behind SSOGs. This leaves the native processes of sequence divergence, overprinting, and remodeling as the next best candidates to explain SSOG origination.

Our analyses go deeper than previous efforts to understand the origin of SSOGs in prokaryotes. Our results are consistent with a previous study where a low fraction of viral matches have been found(Yin & Fischer 2006). In contrast, earlier inquiries into the origin of bacterial orphans from Daubin and Ochman(Daubin & Ochman 2004b, 2004a) investigated compositional similarity between orphans in *E. coli* and bacteriophage sequences known at the time and proposed acquisition from bacteriophages as a reasonable explanation for orphan origin. Our analysis of some of the same features (GC content, CpG content), across thousands of species, for which data was not available at the time, shows instead that orphans closely match their native genomes’ composition. Cortez et al.(Cortez et al. 2009) proposed in 2009 that integrative elements are the major source of ORFans in an analysis of 119 prokaryotic genomes, based on their identification of clusters of genes of atypical composition. But their evidence is tangential and relies on the fact that some of the identified ORFans (39% of 8,428 genes) are found in such clusters of genes with atypical composition, likely to be derived from integrative elements. Furthermore, results on the crucial point of the composition of ORFans themselves are missing from this study. As there is little overlap between the 119 species surveyed by Cortez et al. (entire prokaryotic diversity) and here (>4,000 species of the human gut microbiome), it might be the case that external origination of orphans is more prevalent in species found in different environments.

If most of SSOGs found here were indeed “grown locally”, by what mechanisms did that happen? Extensive sequence divergence is surely one of them. In a previous study, extensive remote homology searches of novel genes in metagenomes using Hidden Markov Model profiles retrieved significant similarity for ∼15% of them(Lobb et al. 2015). Even more sensitive methods might reveal the true percentage to be even higher. In eukaryotes, different approaches have found that many, perhaps most genes without similarity might have simply diverged beyond recognition(Vakirlis, Carvunis, et al. 2020; Weisman et al. 2020). While low syntenic conservation prohibits meaningful application of this method to our dataset, our example in Figure 6C and D clearly shows that syntenic conservation can occur. Divergence of gene duplicates within the same genome can result in additional SSOGs, either by repurposing the same reading frame, or alternative ones. The percentage of SSOGs sharing similarity to conserved genes from the same genome that we report here is likely an underestimate, as divergence beyond detectable similarity is potentially at play at this level as well. A crucial question, outside the scope of this study, is how often this absence of statistically significant similarity should be taken as absence of functional similarity. The alternatives to divergence are de novo gene birth and overprinting. Have these mechanisms been neglected in prokaryotes? In our view, at least within the environment of the human gut, this seems to be the case. Notably, the human gut environment is well-sampled and allows us to detect SSOGs with confidence. Nevertheless, this does not give us a good representation of the entire prokaryotic diversity. For example, gut bacteria are generally anaerobic and strongly associated with the eukaryotic hosts. Thus, their ecology is very different from environmental bacteria and the extent to which our findings generalize to other environments is currently unclear. Future studies that leverage the ever-increasing global metagenomic sequences will be useful to tackle that question. Additionally, experimental evidence shows that the emergence of functional proteins via de novo gene evolution from noncoding sequences can provide adaptation in *Escherichia coli*(Babina et al. 2023; Frumkin & Laub 2023). Thus, de novo evolution may be more plausible, and prevalent, than usually assumed.

A mostly native SSOG origin does not diminish the role of HGT in prokaryotic evolution. Indeed, when one looks at how the percentage of viral matches increases as the number of prokaryotic species in a family increases (Figure 4), it is evident that novel genes can spread between phages and bacteria. It has been known for a long time that phages also invent novel genes(Fremin et al. 2022; Pavesi et al. 2013; Sabath et al. 2012). An intriguing question for future studies then is whether there is any qualitative or quantitative difference between the processes that surely occur in bacteria and phages, or whether both simply serve as input to a common pool of novel sequences that is accessible via horizontal gene transfer and is maintained under selective pressures or neutrally(Wolf et al. 2016; Iyengar & Bornberg-Bauer 2023).

Another important but difficult question is whether our estimates of the number of SSOGs is a good approximation of the true number of entirely novel, *functional* coding sequences unique to each species. Given that SSOGs are overwhelmingly short, it is reasonable to assume that some of them are prediction artefacts(Yu & Stoltzfus 2012). This is an important limitation and we have taken several steps to mitigate it: We have relied on preexisting annotation consistent for all genomes. Note that using consistent, robust annotations is an important measure against spurious results when it comes to estimating species-specific genes(Weisman et al. 2022). Furthermore, we have considered any sequence with an EggNog match as conserved, but it is possible that an evolutionarily constrained gene might converge towards an existing domain or motif. Finally, we have excluded any recently emerged gene from SSOGs that was transferred to another species shortly after having emerged. Notably, novel small ORFs and overlapping reading frames have been omitted in the automatic annotation. Nevertheless, it is plausible that some homologues of the shorter SSOGs could exist in other species but might have been missed by the annotation pipeline due to their short length, which would push SSOG numbers higher. For all these reasons, the presented SSOGs are expected to contain both false positives and false negatives.

Additionally, one can assume that such mis-assignments are even higher for metagenome-assembled genomes (MAGs) which also contribute to the UHGP. First, assembly and binning methods might fail to reconstruct particular species, especially when they are highly diverse. Such missing species could result in false positives orphans in related species. Nevertheless, on average 86% of the reads from a human gut metagenome map to the genomes that are the raw material of the UHGP(Almeida et al. 2021), suggesting that the species diversity in the human gut is well covered. Second, errors at the binning step could lead to the inclusion of wrong genes into MAGs, which then appear to be SSOGs. To mitigate this effect, we have excluded contigs without conserved genes. Third, MAGs can be incomplete and are known to be biased against mobile elements(Maguire et al. 2020). Note that completeness and contamination cutoffs have been applied to the MAGs included in the UHGP(Almeida et al. 2021); however these estimates are based on marker genes and can underestimate the actual values(Meziti et al. 2021; Chen et al. 2020). The bias against mobile elements particularly impacts the accessory genome, where also SSOGs can be found, potentially leading to a bias against SSOGs. Fourth, misassembly or incomplete contigs can lead to the prediction of spurious ORFs. We examined the impact of this bias by comparing the SSOG numbers for species with different assembly qualities and cannot find an association (Supplementary figure 1C). Taken together, the estimated SSOG numbers do not seem to be driven by the inclusion of MAGs in the data set.

When looking for signatures of protein-coding selection at the intraspecies level we found that SSOGs lie on average between noncoding negative controls and conserved positive ones, while largely overlapping both. To be as stringent as possible, we use dN/dS to define a high-confidence SSOG set yet we must stress that these dN/dS values and their comparisons should not be used to definitively accept or reject the functional status of a given SSOG. The reasons, as already discussed, are the technical limitations of dN/dS at the intraspecific level(Kryazhimskiy & Plotkin 2008), and that a recently originated gene can be functional without any selection signatures(Vakirlis et al. 2022; Wacholder et al. 2023) since there might not have been time to accumulate enough mutations for selection to be detectable. What’s more, intraspecific alignments of conserved genes have on average four times as many sequences than those of SSOGs, increasing the statistical resolution and leading to better detection. They are also guaranteed to contain cases of HGTs which would also bias dN/dS values towards those of inter-specific comparisons. Furthermore, it is difficult to extrapolate the percentage calculated among the small subset of SSOGs with enough genetic diversity to all the SSOGs, given that it leads to a bias: SSOGs that have existed long enough to have the necessary diversity are more likely to be functional than those restricted to only one genome. Yet, conservation across multiple genomes, or even species is not always infallible: for example, conserved gene families have been built which were found to be composed of spurious open reading frames (ORFs) located on the noncoding strand of another conserved gene family(Eberhardt et al. 2012). Nevertheless, functional evidence for such “antisense proteins” shows that the noncoding strand also provides potential for evolutionary innovation(Ardern et al. 2020), making it difficult to distinguish functional and non-functional antisense proteins.

Our estimate of the number of SSOGs should be viewed with caution and with all the aforementioned caveats in mind. However, the primary focus of our study is not the number of SSOGs per se, but rather what we can glean about their origins when subjecting them to comparisons. Notably, using the dN/dS defined high-confidence set or a more stringent length cut-off does not impact our main conclusion demonstrating that our results are not driven by false positive SSOGs. In any case, future studies might provide further experimental insights into the expression and function of the SSOGs described here. Approaches incorporating transcriptomics, ribosome profiling, and high-throughput functional assays might be especially informative. To distinguish functional expression from pervasive transcription and translation, analysis of differential expression of SSOGs under different conditions is of particular interest.

We have identified more than half a million SSOGs and even if a large part of them turns out to be spurious, we would still be left with thousands of candidates that could be expressed as entirely novel proteins within the boundaries of the human body. It is more and more acknowledged that pangenome diversity is important in the human gut microbiome and might have consequences for human health(Vatanen et al. 2019). SSOGs within strains of human gut bacteria could perhaps interact with the host, contributing to immune or inflammatory responses and potentially even disease.

While such a link of prokaryotic SSOGs to human health can only be speculative, their overall importance for niche-specific adaptations is far from it. Although the cloud of rare genes has been found to be an integral part of all pangenomes, the evolutionary processes that generate and maintain cloud genes are not well understood and explanations range from niche-specific genes to a constant inflow of transient genes(Baumdicker & Kupczok 2023). Particularly, a class of transient genes with effectively instantaneous gene replacement rates has been suggested to contribute to the cloud fraction of microbial pangenomes(Wolf et al. 2016). Here we highlight that SSOGs are an important component of prokaryotic pangenomes, and especially of the cloud fraction, suggesting that they are mostly transient. Thus, most SSOGs might be removed quickly after their origination, where some of them could prove adaptive and persist in the population, resulting in non-cloud SSOGs and – after longer evolutionary time – even in conserved genes. To gain a comprehensive picture of prokaryotic evolution, we thus need to understand where SSOGs come from, what specific functions they assume and how they evolve to be maintained in the population.

## Materials and Methods

### SSOG identification

Species-specific orphan genes (SSOGs) were identified as follows: Initially, the UHGP50 v.1 catalogue which is clustered into protein families at 50% identity by Almeida et al.(Almeida et al. 2021) was parsed and families with members in more than one species were removed. Next, we removed families with a precomputed EggNOG(Hernández-Plaza et al. 2023) match (based on Almeida et al. data for their representative sequence, i.e. the first in order sequence in the family file). We then performed two similarity searches against the entire NCBI prokaryotic RefSeq database (downloaded January 2022), using DIAMOND(Buchfink et al. 2021) v2.0 *blastp* with the representative protein sequence of each family as query: first using DIAMOND “fast” mode, and removing any sequence that had a significant match (E-value<0.001), except for sequences with >90% coverage and >95% identity that can reasonably be expected to be derived from the same species as the query sequence, present in RefSeq. The same search was then performed for the remaining sequences using the “ultra-sensitive” mode and the same criteria were applied. A final search was performed using BLASTp(Altschul et al. 1997) against the entire UHGP50 protein catalogue (consisting of the representative protein sequences) with an E-value cut-off of 10^−5^ and “*-max_target_seqs 1000”* and any candidate SSOG sequence matching with a protein from a different species was filtered out. The remaining sequences constitute the set of SSOGs. To mitigate contamination, we also filtered out candidate SSOGs found on contigs that only contain SSOGs (only the representative sequence was tested for this). All the non-SSOG families (i.e. those removed in the previous steps) were henceforth considered “conserved” and we counted the number of unique species represented within it.

### Prediction of CDS and protein properties

Protein secondary structure and intrinsically disordered regions were predicted using the RaptorX Predict_Property package v1.01(Wang et al. 2016) in default mode. Protein self-aggregation was predicted using PASTA(Walsh et al. 2014) v2.0 in default mode with an energy threshold of −5, and the percentage of the protein expected to form aggregate-prone regions was used. Transmembrane domains were predicted using Phobius(Käll et al. 2007). Codon Adaptation Index (CAI) was calculated for all sequences using the CAIJava(Carbone et al. 2003) tool with arguments *“-s -i 15 -k 3 -g”*, and for *E. coli* sequences additionally using the *codonw*([CSL STYLE ERROR: reference with no printed form.]) tool and the EMBOSS *cai* tool, both using their respective *E. coli* set of reference genes. GC% and GC% in the 3^rd^ synonymous codon position was calculated with *codonw*([CSL STYLE ERROR: reference with no printed form.]). CpG dinucleotide frequency was calculated in *R* with the SeqinR(Charif & Lobry 2007) package. Biosynthetic cost was calculated by averaging the Akashi and Gojobori(Akashi & Gojobori 2002; Barton et al. 2010) amino-acid scores for each protein sequence. Isoelectric point was calculated with the *R* package *peptides*(Osorio et al. 2015).

### Similarity searches

We downloaded two gut phage protein sequence databases in FASTA format, MGV(Nayfach, Páez-Espino, et al. 2021) and GPD(Camarillo-Guerrero et al. 2021). Additionally, protein sequences were extracted from phage genomes assembled by Shah et al.(Shah et al. 2023) and downloaded from http://copsac.com/earlyvir/f1y/gbks/ in May 2022. Similarity searches against viral protein databases were conducted using DIAMOND *blastp* in ‘very sensitive’ mode. The criteria applied to define significant hits, in increasing order of stringency, were the following: identity <95% & E-value < 10^−3^ (low), identity <95% & E-value < 10^−5^ (medium), 95%>identity >40% & query coverage % >40% & E-value < 10^−5^ (high), 95% > identity > 60% & query coverage >70% & E-value < 10^−5^ (very high). To identify SSOGs potentially derived from conserved native genes, we also conducted similarity searches of SSOGs against all annotated CDS in their respective genomes using NCBI’s TBLASTX with 80% identity, 70% query coverage and E-value 10^−5^.

### Genomic and functional features

We predicted the presence of prophages in all 286,997 prokaryotic genomes by running Phigaro(Starikova et al. 2020) in default mode. We then counted the number of SSOGs and non-SSOGs that are situated within the predicted prophage genomes.

All general genomic data such as assembly quality, as well as taxonomic information were retrieved from the ‘genomes-all_metadata.tsv’ table provided by Almeida et al. The species-level phylogenetic tree used was taken from the file ‘bac120_iqtree.nwk’ provided by the authors. All pangenome related statistics were calculated using the pangenome files already available from UHGP, and accessed through links such as ftp://ftp.ebi.ac.uk/pub/databases/metagenomics/mgnify_genomes/human-gut/v1.0/uhgg_catalogue/MGYG-HGUT-000/MGYG-HGUT-00001/pan-genome/ where “MGYG-HGUT-00001” is a species code. Protein/CDS IDs (e.g. GUT_GENOME255189_00514) were matched to unique gene locus IDs (e.g. epsF_2) used within pan-genomes by Almeida et al. and then statistics such as number of genomes where a gene is present were extracted by the relevant tables provided by the authors (‘genes_presence-absence_locus.csv’ files). The categorization of core (defined at 90% presence) and accessory genes was also retrieved from these directories (‘accessory_genes.faa’ files). To count the number of SSOGs per pangenome, we counted the number of unique gene locus IDs, in each species, that we had classified as SSOGs (one name identifies a single locus and is present only once, even if found in multiple genomes; provided by Almeida et al.).

We sought to identify SSOGs among those prokaryotic genes included in the fitness browser(Price et al. 2018). We downloaded protein sequences of fitness browser proteins as well as raw t-score tables for each species from the fitness browser website. We then used DIAMOND to search for similarity between our SSOG proteins and those fitness browser proteins. We considered a hit as significant if it had E<10^−5^, % identity > 70%, and 80% query coverage. We then filtered matches to retain only those to fitness browser proteins with statistically significant fitness effects, defined as t-score > 4 in at least one condition (as detailed in the original publication). To only consider unambiguous cases, we looked only for SSOGs matching to a unique fitness browser protein.

### Identification of orthologous regions and definition of de novo candidates

For each SSOG, we searched for each orthologous region in two sets of genomes. In every genome of the same species without the SSOG, and in every genome of its closest neighboring species, as defined by the species-level phylogeny provided by Almeida et al(Almeida et al. 2021). To identify the orthologous region in each genome, we took the immediate upstream and downstream SSOG neighbors and searched whether a homologue, that is, a member of the same family, exists in that genome. We only considered cases where a single homologue existed, that is, cases with paralogues were discarded. If both a homologue of the downstream and the upstream neighbor existed in a given outgroup genome, and these were separated by at most 1 gene, we considered the region as conserved in synteny. In order for a SSOG to be characterized as a de novo candidate, such a conserved syntenic region had to be identified in at least one genome of the same species and at least one genome from the closest species. Additionally, we discarded regions where an unannotated ORF with at least 80% query coverage and 60% similarity to the SSOG existed in the syntenic region of the outgroup species. Finally, we discarded 23 candidates which had similarity matches to conserved genes from the same genome (see ‘Similarity searches’ subsection above).

### Identification of operon-like genomic regions

To identify operon-like gene arrangements, we scanned all UHGG genomes and took advantage of the existing Gene Ontology (GO) annotations available from the InterPro searches performed by the authors. For every gene of every genome, we examined its three immediate downstream and three immediate upstream neighbors. If all seven genes were on the same contig, had the same orientation, and at least three of the neighbors shared at least one GO term annotation, then that gene was considered to be in an operon-like arrangement, and the GO term was associated to that gene. Note that we merged the terms GO:0016020 and GO:16021 to be consistent with the current version of GO, after we noticed that terms related to the membrane that came up later in our analyses had been tagged as obsolete and collapsed into the general term “membrane”. All associations (one gene can have more than one if more than one GO term is found more than three times) for all genes were formatted into a custom annotation file. This file was used as the background annotation, together with a list of SSOGs in BiNGO v3.0.3(Maere et al. 2005) to perform the overrepresentation analysis. The options used were: hypergeometric test, Benjamini-Hochberg FDR correction and a 0.05 threshold. For analysis restricted to each taxonomic class, the same was performed but only including genes from species of a given class.

In the case of SSOG GUT_GENOME108422_00105 (representative of a family of 135 proteins) our analysis of the syntenic region showed the gene to be present at the exact orthologous locus as SSOG GUT_GENOME123088_00522 (representative of a family of 866 members) both belonging to species PeH17 (GUT_GENOME014725). Aligning the two representative sequences gave results consistent with the criteria used by Almeida et al. for protein clustering: the sequences align with 100% identity but only over their first 25 residues (46% of the length of the shortest sequence), while the rest of the sequences align very poorly. However, all other protein sequences of the GUT_GENOME123088_00522 family align nearly perfectly with any protein from the GUT_GENOME108422_00105 family, with no gaps and with 94% identity. Indeed, aligning all CDS sequences from both families together results in an alignment with average pairwise percent identity of 97% (calculated using the *goalign* tool and its *compute distance* method with the *pdist* metric). Furthermore, a nucleotide alignment of the representative sequences showed that their difference at the protein level is due to a frameshift. Overall, these two families, with the exception of the representative sequence of GUT_GENOME123088_00522, would have been expected to cluster together given the parameters used by Almeida et al. (*linclust* with 80% coverage and 50% identity).

### Selection signatures, quantification and statistical analysis

All statistics were done in R v3.6.2. Plots were generated using *ggplot2*(Wickham 2011). All statistical details including the type of statistical test performed and exact value of n (n represents either number of genomes or number of genes) can be found in the Results and figure legends. Boxplots show median (horizontal line inside the box), first and third quartiles of data (lower and upper hinges) and values no further or lower than 1.5*distance between the first and third quartiles (upper and lower whisker). No methods were used to determine whether the data met assumptions of the statistical approaches. Subsampling of conserved genes to control for length difference to SSOGs was performed with replacement, using a customized version of inverse transform sampling. We sampled conserved genes belonging to each species separately, and only species with at least 100 conserved and at least 10 SSOGs were included (n=3,769). The length median of the sampled conserved genes was 228nt and the mean was 305nt, compared to 225nt and 300nt in SSOGs respectively. To detect signatures of selection acting on protein-coding sequences, we first generated DNA multiple sequence alignment of the CDSs of 154,650 SSOG families with at least 2 non-identical sequences. We then used the *yn00* executable of PAML(Yang 2007) to calculate dN, dS and omega (dN/dS) with the Yang & Nielsen 2000 method(Yang & Nielsen 2000). The program was run with default values and was given as input the aforementioned MSA. The dN, dS (with their estimated standard errors) and dN/dS values were obtained for all pairwise combinations of members of each family. For each family, to only take into account statistically meaningful comparisons we kept only those comparisons were dS>0 and SEdS/dS < 1. We then calculated the mean for a given family from the remaining pairwise comparisons. We also calculated single alignment and tree omega using HyPhy(Kosakovsky Pond et al. 2020) under the MG94xREV model (using the steps provided by the authors here: https://stevenweaver.github.io/hyphy-site/tutorials/current-release-tutorial/#estimate-a-single-alignment-wide, but without optimizing the branch lengths) and a phylogenetic tree generated with a “quick-and-dirty” analysis by RAxML next generation(Kozlov et al. 2019) (*raxml-ng*) with the following command: “*raxml-ng --msa {INPUT ALIGNMENT} --model GTR+G --search1*”. The same analyses (avg. pairwise dN/dS with PAML and omega with HyPhy) were performed on three negative controls based on the initial SSOG MSA: 1) alignments frame-shifted by one nucleotide (i.e. on the +1 frame) using *goalign*(Lemoine & Gascuel 2021) (*goalign trim -s -n 1* followed by *goalign trim -n* 2) and for which in frame stop codons were removed 2) alignments where entire alignment columns were randomly shuffled using the Multiperm(Anandam et al. 2009) tool with the option -- conservation=none and 3) completely randomized alignments where each sequence was randomly shuffled separately. The conserved, positive controls were generated as follows. We selected randomly one species from each of the nine best represented taxonomic classes, that had between 100 and 200 available genomes, so as to have enough sequences in the alignments while keeping the analysis tractable. The following species were selected: GUT_GENOME000122, GUT_GENOME001519, GUT_GENOME113559, GUT_GENOME001122, GUT_GENOME286320, GUT_GENOME147777, GUT_GENOME220136, GUT_GENOME096385, GUT_GENOME009256. Available genomes were downloaded from UHGG v1. In the genomes of each species, we retrieved genes with known assigned names (e.g. mtaB), which correspond to conserved genes with known functions. Genes were grouped based on their exact name. The sequences of each gene were then aligned in a codon-aware manner using *codonalign* from the *goalign* tool to account for frameshifts. The rest of the analysis was performed as for SSOGs above.

## Acknowledgements

We thank Daniel Tamarit and Franz Baumdicker for valuable comments on an earlier version of the manuscript. We acknowledge funding from the DFG in the framework of the SPP2141 to A.K. (KU3610/2-1). The research project was supported by the Hellenic Foundation for Research and Innovation (H.F.R.I.) under the “3rd Call for H.F.R.I. Research Projects to support Post-Doctoral Researchers” to N.V. (Project Number:7330). This research was supported in part through high-performance computing resources available at the Kiel University Computing Centre.

## Contributions

AK and NV conceived the study. AK and NV wrote the manuscript. AK supervised the study. NV performed the analyses.

