## Supplementary figures and supplementary table legends for "Large-scale investigation of species-specific orphan genes in the human gut microbiome elucidates their evolutionary origins"

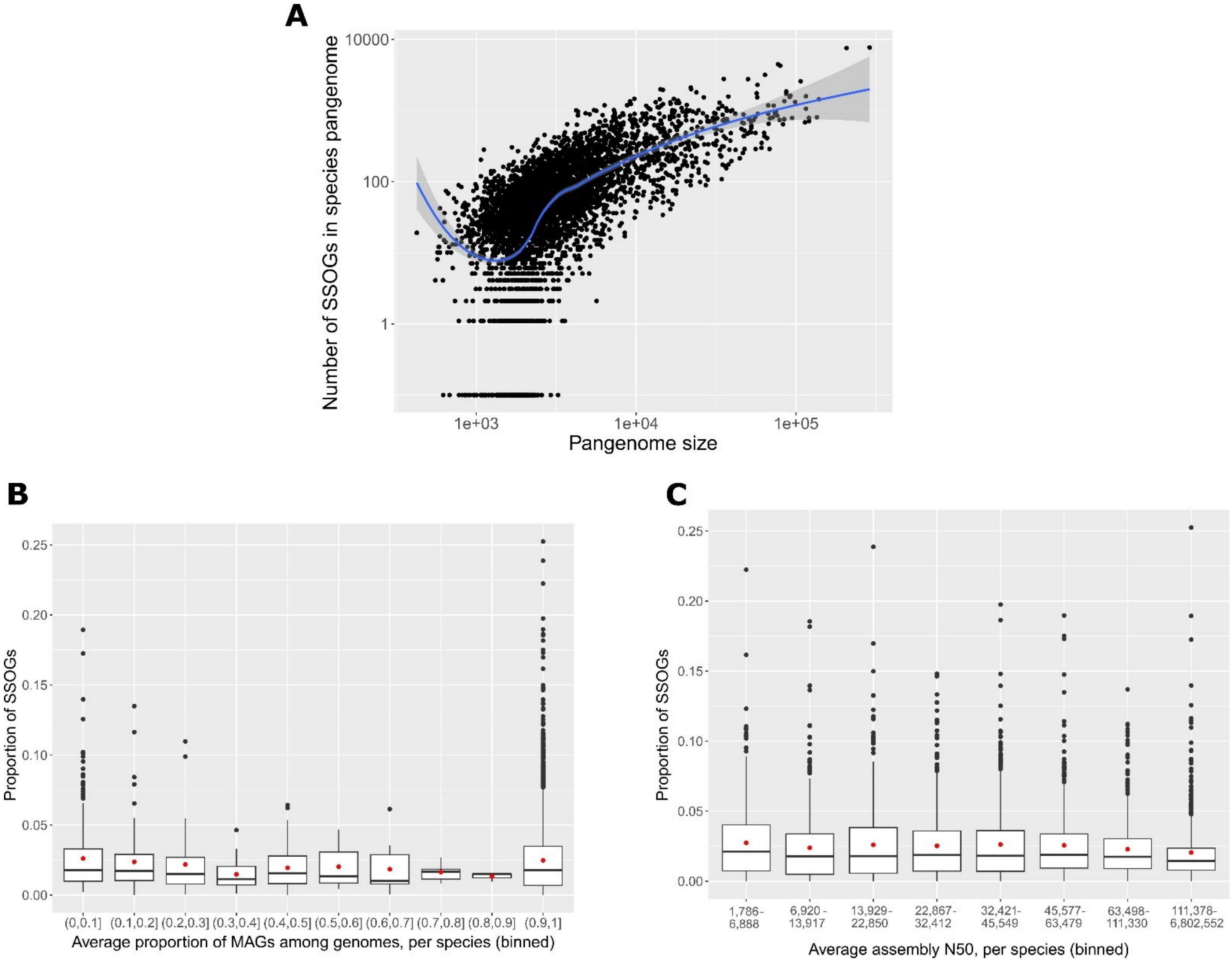


**Supplementary Figure 1**. **A**: Number of SSOGs in species pangenome as a function of pangenome size. Data are log-transformed. A LOESS fit in the log-transformed data is shown. We note that some saturation is visible in values of highest pangenome sizes. **B**: Proportion of SSOGs as a fraction of the pangenome, in species binned according to their MAG proportion. **C**: Same as B, but species are binned according to their average assembly quality N50. Bins have been set to contain the same number of SSOGs.

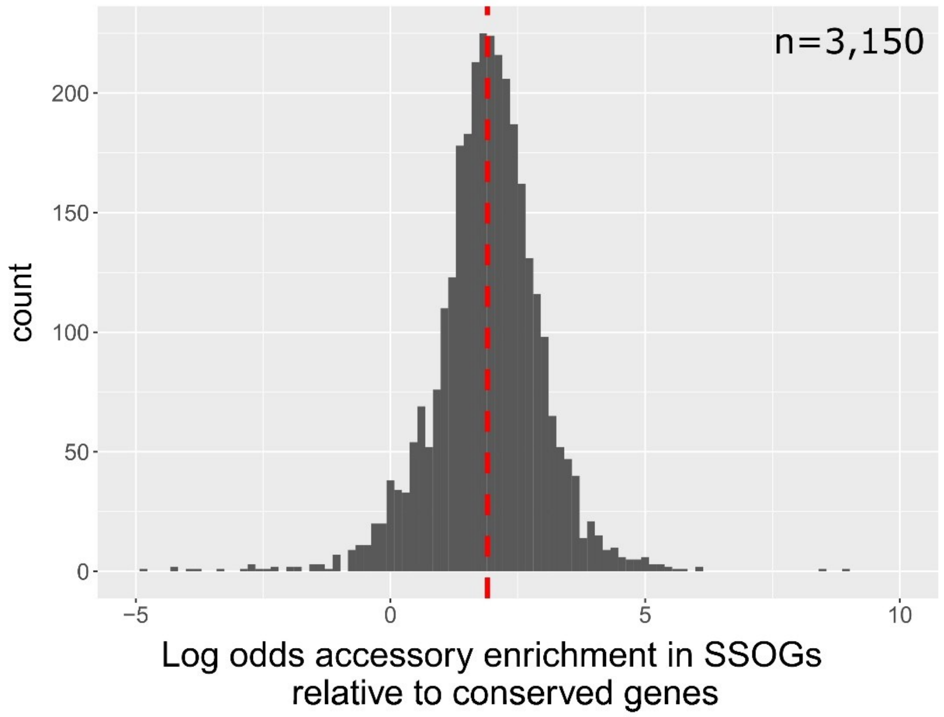


**Supplementary Figure 2**: Enrichment of accessory genes over core genes in SSOGs relative to conserved genes (see Methods). Range of X-axis has been reduced for visibility, 12 genomes with extreme values are not shown. Core and accessory genes were defined by Almeida et al. (doi.org/ 10.1038/s41587-020-0603-3).

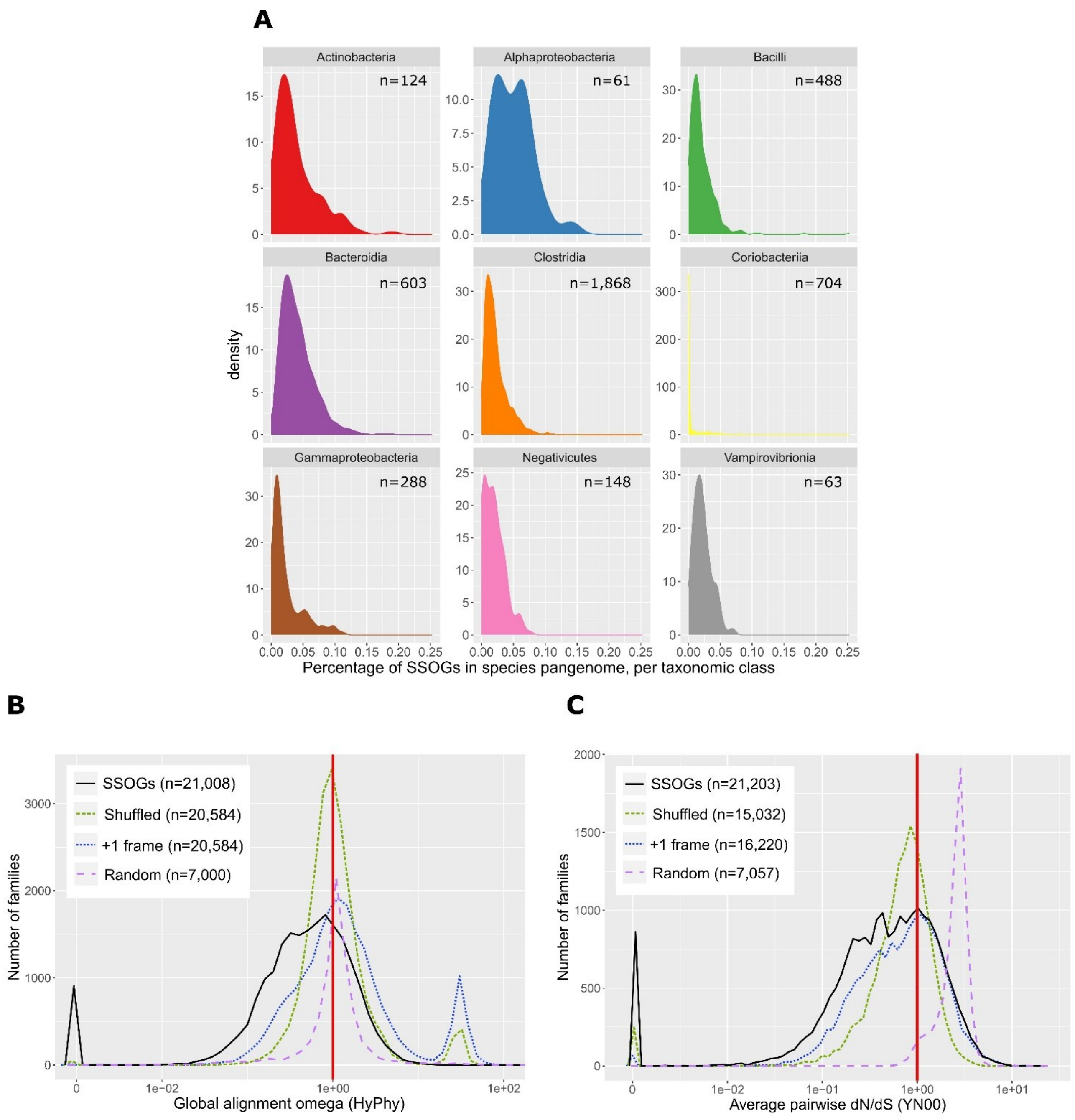


**Supplementary Figure 3. A**: Distributions of percentage of SSOGs in each species pangenome split across the nine best represented taxonomic classes. **B**: Distributions (60 bins) of global alignment omega calculated using HyPhy, for all SSOG families meeting our genetic diversity criteria and containing at least 4 sequences, and a set of negative controls (see Methods). Red line marks omega=1 which corresponds to neutral evolution. Four extreme positive outlier values over 100 are not included for visibility. **C**: Distributions (bins=60) of average pairwise omega (dN/dS) per family for SSOGs with CDS length > 300nt (solid black line) and three matching negative controls (see Methods). Red line marks dN/dS=1 which corresponds to neutral evolution.

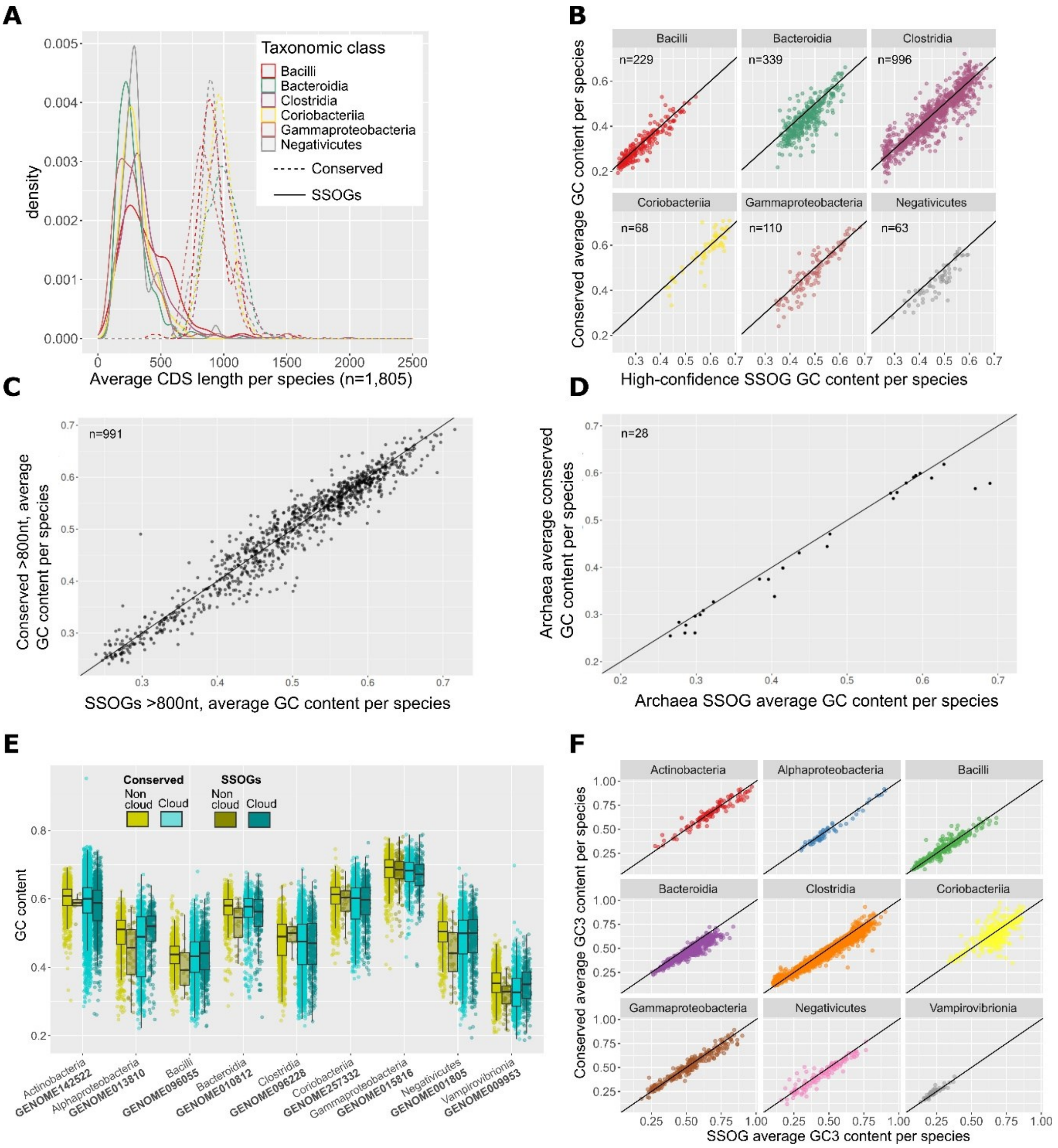


**Supplementary Figure 4**: **A:** Density plots of the average length of high-confidence SSOGs and conserved CDSs in each species, grouped by taxonomic class. **B:** Average GC content of high-confidence SSOGs and conserved genes in each species, grouped by taxonomic class. **C:** Average GC content of SSOGs and conserved genes longer than 800nt in each species. **D:** Average GC content of SSOGs and conserved genes in Archaea species. **E:** Intraspecific comparison of GC content in SSOGs and conserved genes that are found in 10% or less of genomes in their species (cloud) or not (non-cloud). Data are shown for a sample of 10 species with at least 200 SSOGs and at least 10 genomes, drawn randomly from each of the nine best represented taxonomic classes. Each point represents one gene, that is, data here are not averaged as in Figure 3. **F**: Comparison of per species average GC3s between SSOGs and conserved genes in the nine best represented taxonomic classes.

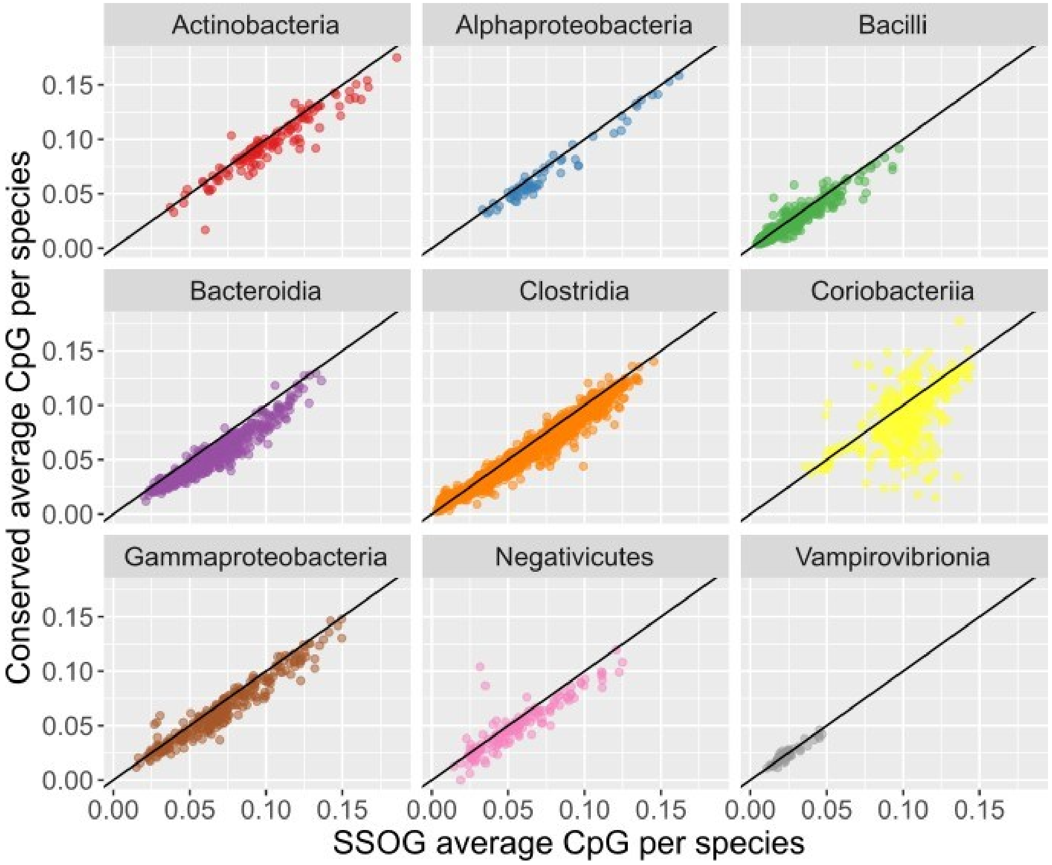


**Supplementary Figure 5**: Comparison of CpG between SSOGs and conserved genes in the nine best represented taxonomic classes.

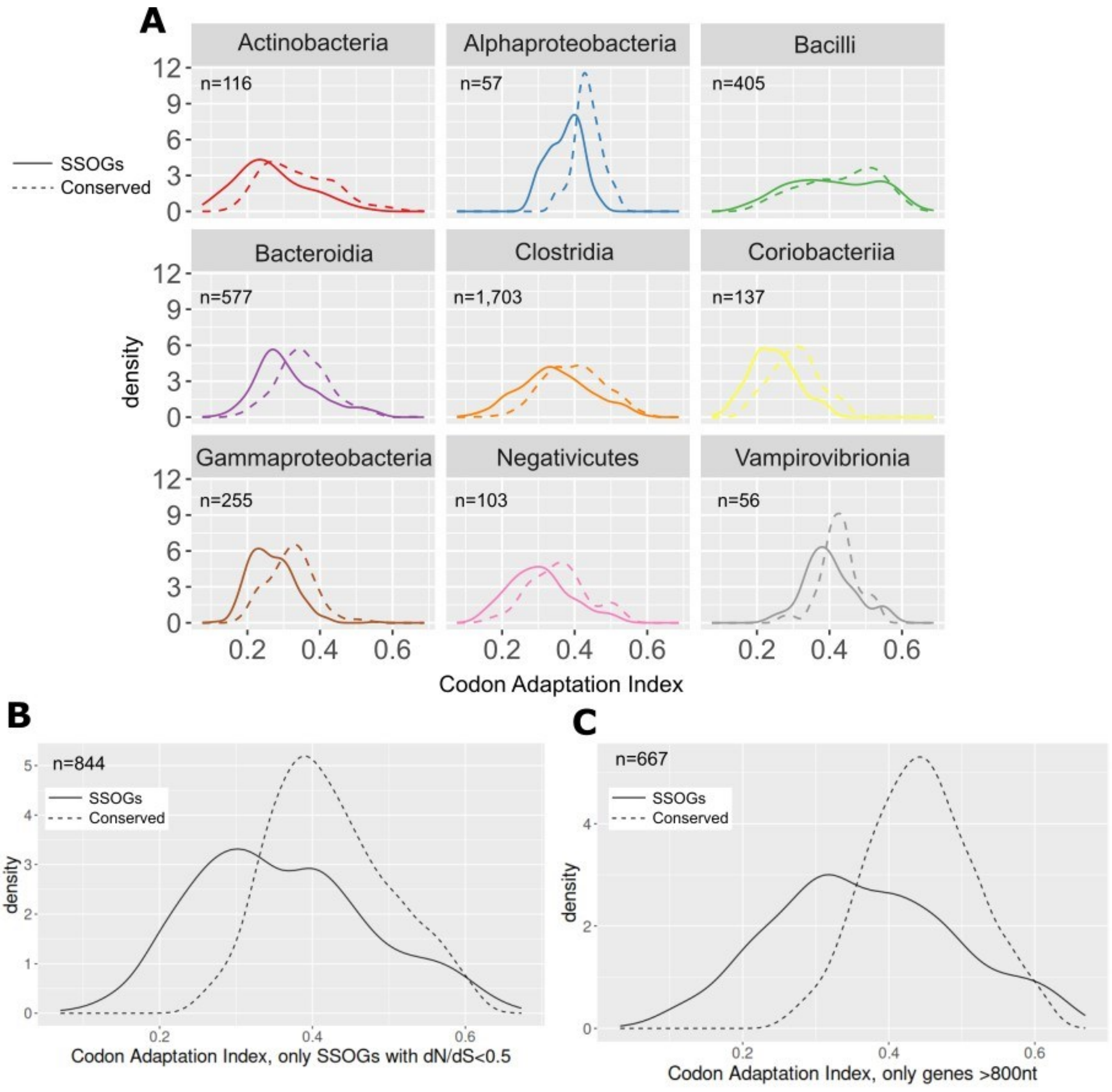


**Supplementary Figure 6: A**: Comparison of distributions of per-species average CAI values between SSOGs (solid line) and conserved genes (dotted line), subsampling conserved genes to match the length distribution of SSOGs. All differences are statistically significant with Wilcoxon test P-value < $10^{-4}$ except for Bacilli (P=0.02) and Vampirovibrionia (P=0.005). **B:** Comparison of distributions of per-species average CAI values between SSOGs (solid line) and conserved genes (dotted line), for the high-confidence set of SSOGs (dN/dS<0.5). **C:** Comparison of distributions of per-species average CAI values between SSOGs (solid line) and conserved genes (dotted line), when taking into account only genes >800nt.

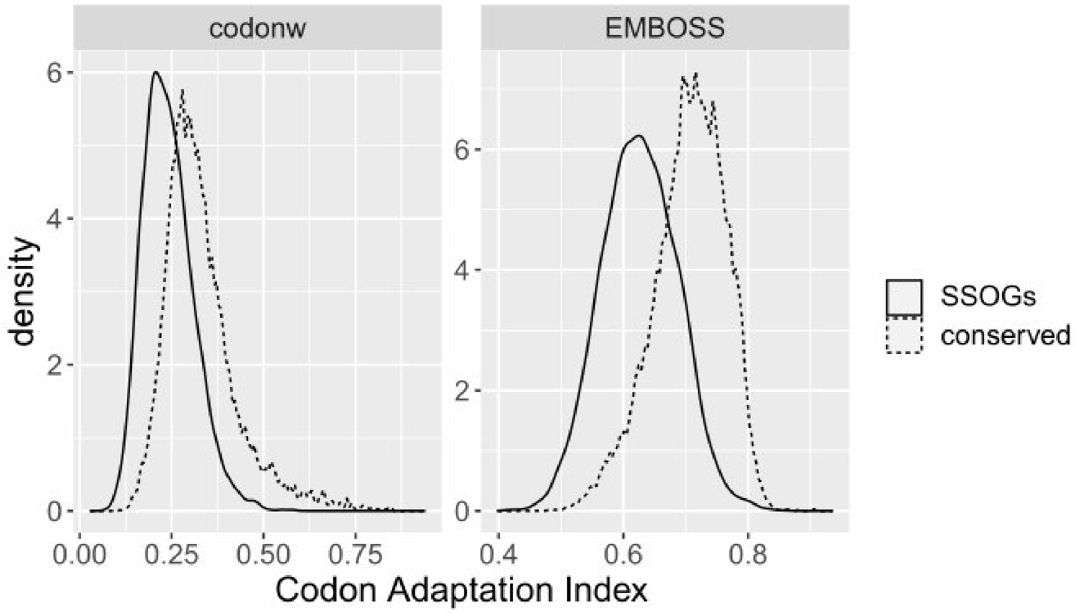


**Supplementary Figure 7**: CAI comparison of all SSOGs (n=4,703) and all conserved genes (n=35,300,683) from 8,288 *E. coli* genomes included in our dataset, calculated using two different tools/reference gene sets. Both Wilcoxon test P-values $< 2.2*10^{-16}$.

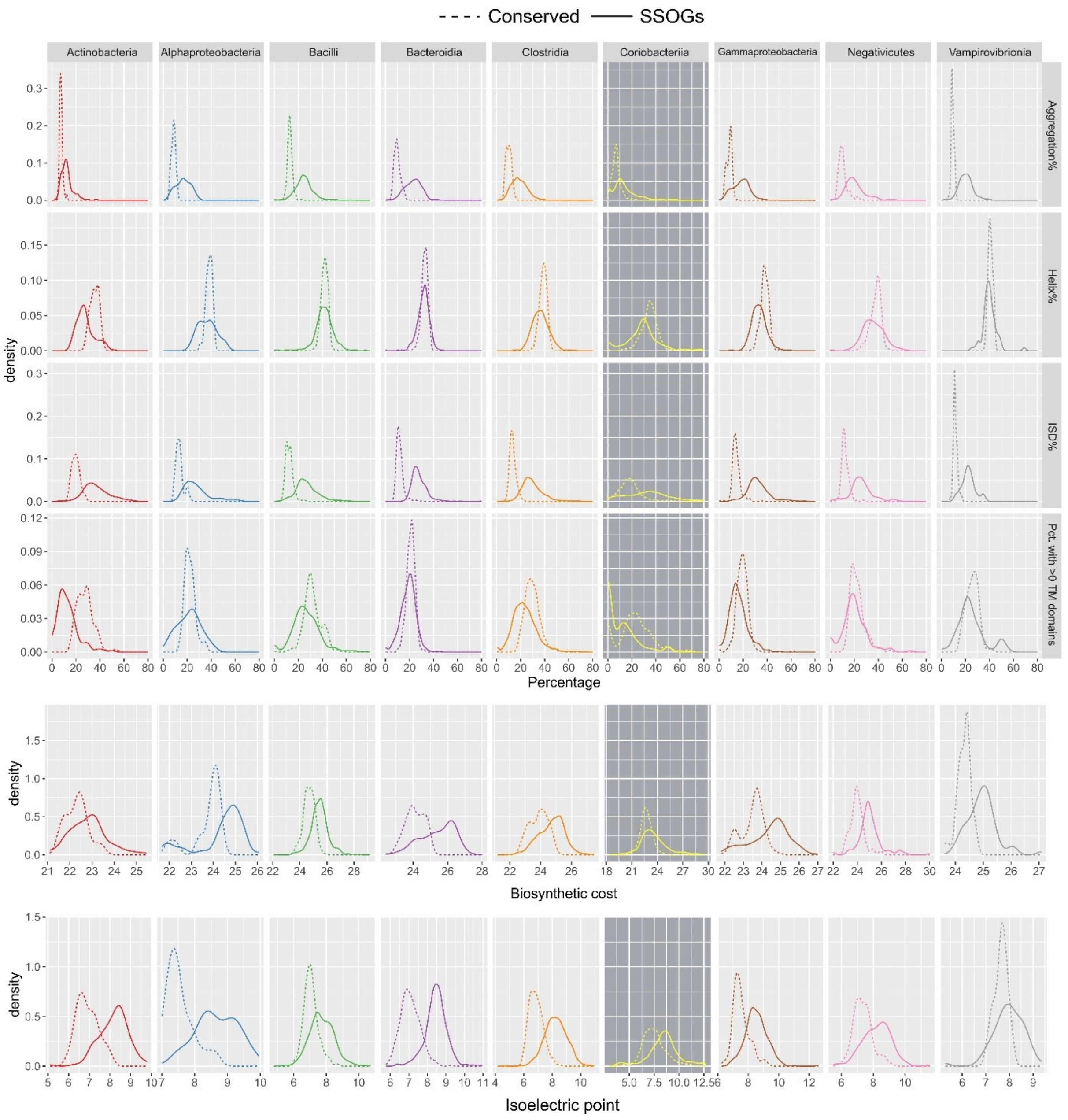


**Supplementary Figure 8**. Comparison of proteins encoded by conserved genes and SSOGs, in each taxonomic class, in terms of average percentage of proteins predicted to self-aggregate, percentage of protein predicted to be helical, percentage of protein predicted to be disordered, percentage of proteins with at least 1 transmembrane domain, average isoelectric point and average biosynthetic cost.

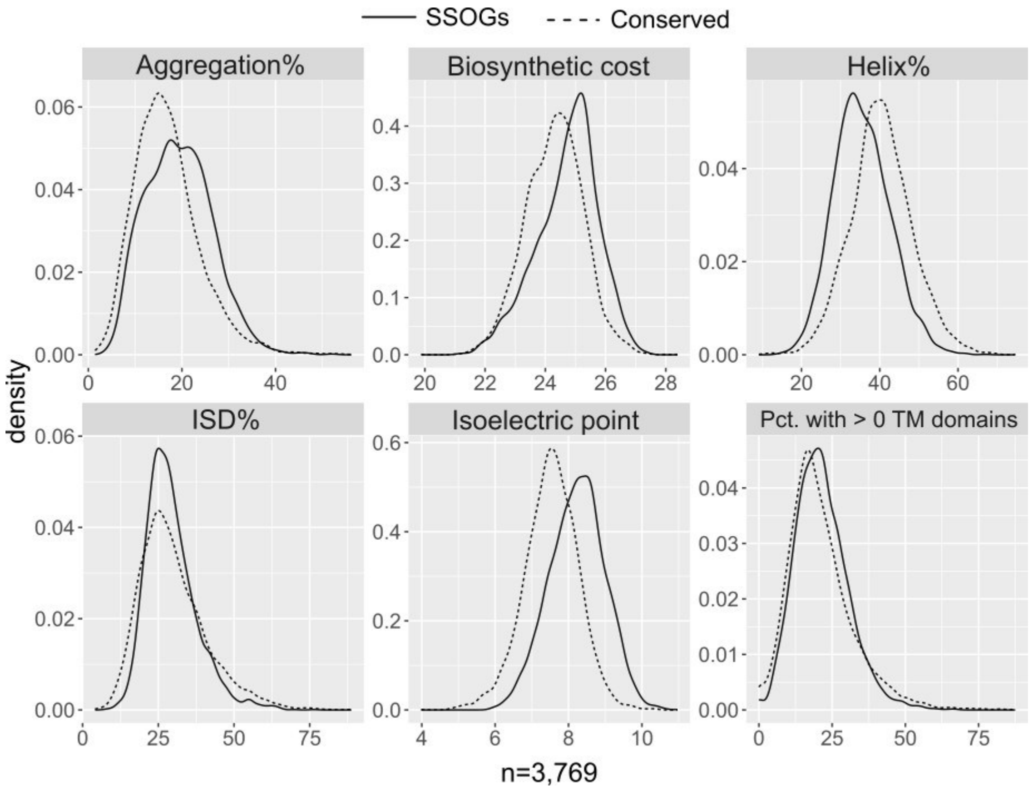


**Supplementary Figure 9**. Comparison of proteins, where conserved proteins have been subsampled in each species to have similar average lengths as SSOG proteins. Comparison of proteins in terms of average percentage of protein predicted to self-aggregate, percentage of protein predicted to be helical, percentage of protein predicted to be disordered, percentage of proteins with at least one transmembrane domain, isoelectric point and biosynthetic cost. No statistically significant difference exists in intrinsic protein disorder (Wilcoxon test P=0.6). Difference in percentage with at least one transmembrane domain is statistically significant ($P=3.7*10^{-7}$) but with a negligible effect size (Cliff's delta 0.07). All other P-value$<2.2*10^{-16}$

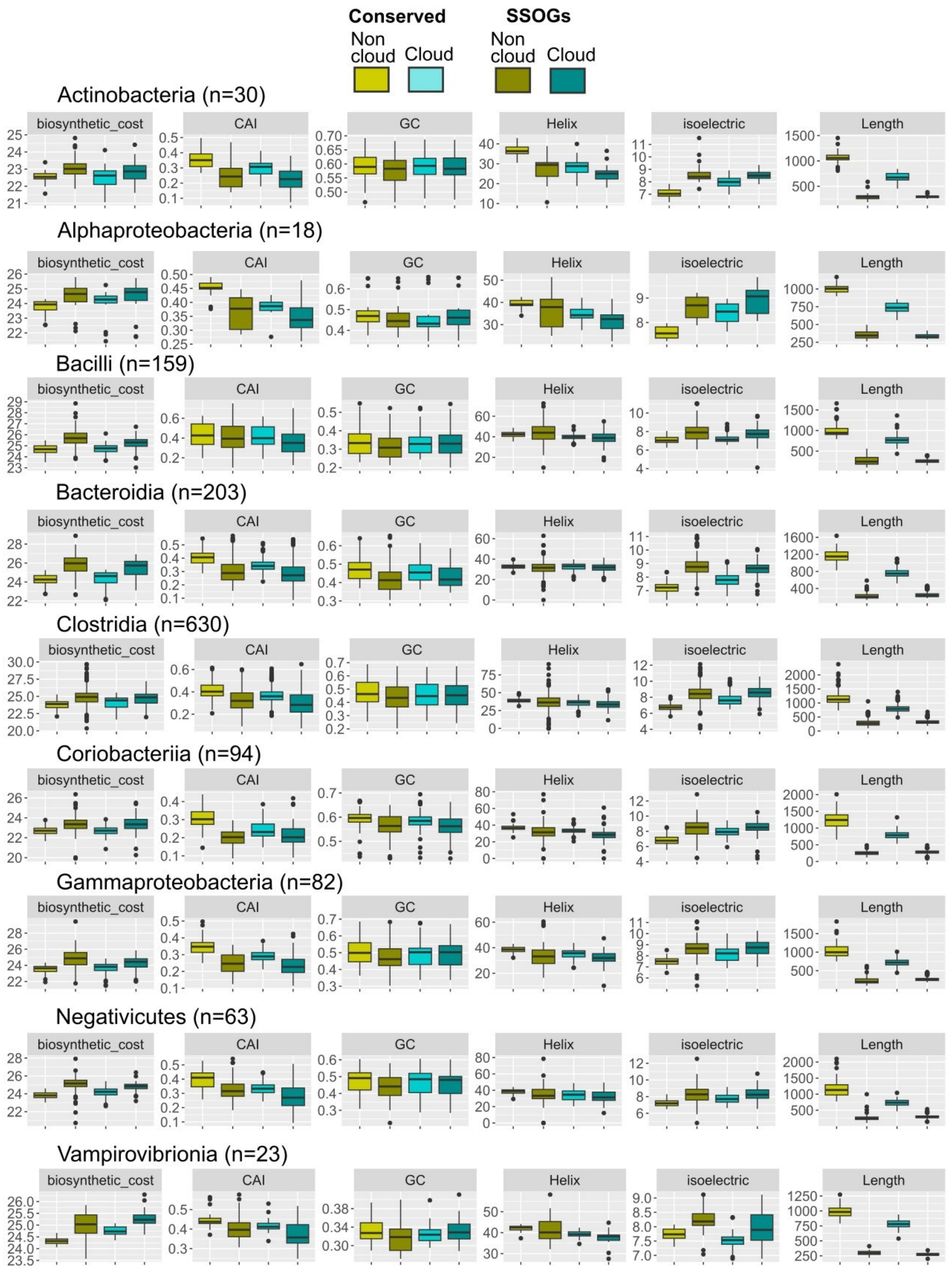


**Supplementary Figure 10.** Comparison of protein biosynthetic cost, CAI, GC content, percentage of protein predicted to be helical, isoelectric point of a protein, and CDS length in 1,302 species belonging to the nine best represented taxonomic classes, and where each species has at least 10 genomes. Data are shown for SSOGs and conserved genes split in cloud (present in 10% of species or less) and non-cloud. Each data point represents an average value per species.

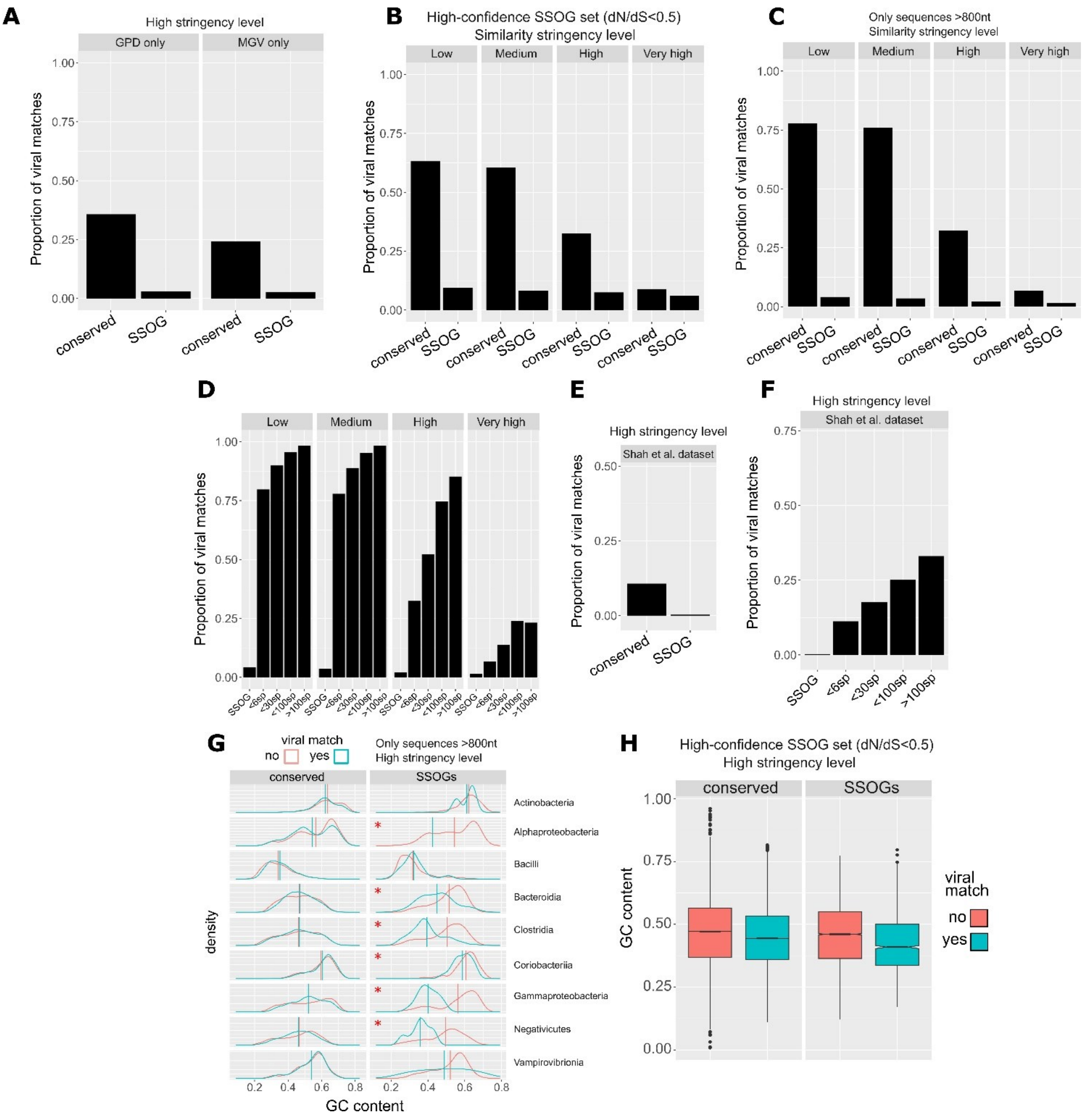


**Supplementary Figure 11. A:** Proportions of SSOGs and conserved genes with viral matches, at high stringency level, in two separate human gut viral protein databases. **B:** Proportions of high-confidence SSOGs with dN/dS<0.5 and conserved genes, with viral protein matches at different levels of stringency. **C:** Proportions of SSOGs and conserved genes longer than 800nt, with viral protein matches at different levels of stringency. **D**: Same proportions as C but including all genes and grouped in bins according to the number of species present. **E**: Same as A but using the Shah et al. viral protein dataset. **F**: Same as D but using the Shah et al. dataset at high stringency. **G**: Differences in GC content between genes longer than 800nt, with and without viral match, in SSOGs and conserved genes. Vertical lines show distribution means. Red asterisk denotes a non-negligible effect size as measured by Cliff's

Delta. **H:** Same as G but for the SSOG high-confidence set. All classes are pooled due to lower sample size.

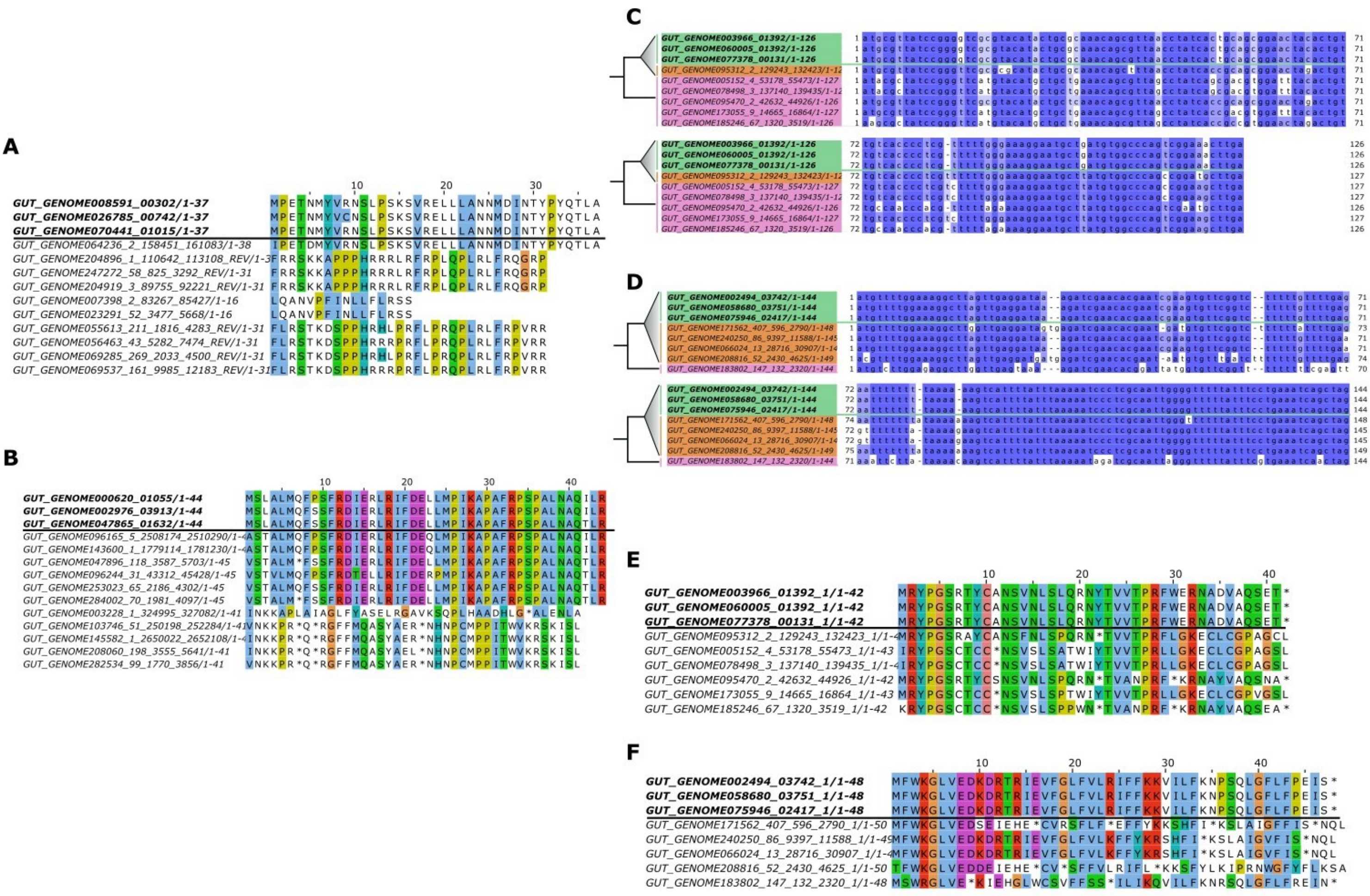


**Supplementary Figure 12. A,B:** Protein translations of the two de novo candidate examples shown in Figure 5 and their orthologous sequences in outgroup genomes. De novo protein sequences shown in bold. Note that the sequences are not aligned. **C**: Alignment of a de novo candidate gene from an unnamed species of genus *Pauljensenia* (green) to its orthologous regions in genomes of the same species that do not have annotated homologues (orange) and genomes of its closest outgroup species (pink). Identical orthologous sequences have been removed from the alignment. **D**: Same as C, but for another candidate from *Enterococcus raffinosus*. **E** : Protein translations of the two de novo candidate examples shown in C and D, and their orthologous sequences in outgroup genomes. De novo protein sequences shown in bold. Note that the sequences are not aligned.

### Supplementary Tables

**Supplementary Table 1**

List of representative de novo candidates.

**Supplementary Table 2**

Results of GO term enrichment analysis over all genes in operon-like arrangements.

**Supplementary Table 3**

Results of GO term enrichment analysis per taxonomic class. Note that rows are not sorted.

**Supplementary Table 4**

Representative Clostridia SSOGs in spore germination-associated operon-like arrangements.
